# Evolution and genetics of accessory gland transcriptome divergence between *Drosophila melanogaster* and D. *simulans*

**DOI:** 10.1101/2023.05.02.539169

**Authors:** Alex C. Majane, Julie M. Cridland, David J. Begun

## Abstract

Studies of allele-specific expression in interspecific hybrids have provided important insights into gene-regulatory divergence and hybrid incompatibilities. Many such investigations in Drosophila have used transcriptome data from whole animals or gonads, however, regulatory divergence may vary widely among species, sex, and tissues. Thus, we lack sufficiently broad sampling of tissues to be confident about the general principles of regulatory divergence. Here we seek to fill some of these gaps in the literature by characterizing regulatory evolution and hybrid misexpression in a somatic male sex organ, the accessory gland, in F1 hybrids between *Drosophila melanogaster* and *D. simulans.* The accessory gland produces seminal fluid proteins, which play an important role in male and female fertility and may be subject to adaptive divergence due to male-male or male-female interactions. We find that *trans* differences are relatively more abundant than *cis*, in contrast to most of the interspecific hybrid literature, though large effect-size *trans* differences are rare. Seminal fluid protein genes have significantly elevated levels of expression divergence and tend to be regulated through both *cis* and *trans* divergence. We find limited misexpression in this organ compared to other Drosophila studies. As in previous studies, male-biased genes are overrepresented among misexpressed genes and are much more likely to be underexpressed. ATAC-Seq data show that chromatin accessibility is correlated with expression differences among species and hybrid allele-specific expression. This work identifies unique regulatory evolution and hybrid misexpression properties of the accessory gland and suggests the importance of tissue-specific allele-specific expression studies.

## INTRODUCTION

Gene expression phenotypes such as transcript abundance, splicing, tissue or developmental stage of expression, and environment-induced plasticity, may be related to many downstream organismal phenotypes and may be acted upon by stabilizing, directional, or diversifying selection. The gene regulatory variants, which may be influenced by selection or drift, and must ultimately explain within and between species expression variation, can be broadly classified into *cis-*acting components, such as promoters and enhancers, and *trans-* acting components, such as transcription factors (Rabinow and Dickinson 1981; Dickinson, Rowan, and Brennan 1984; Patricia J. Wittkopp, Haerum, and Clark 2004; Gibson et al. 2004; P. J. Wittkopp 2005; Ronald et al. 2005; Ronald and Akey 2007). Because the genetic control of gene expression can involve several sites spanning both cis- and trans-acting factors, selection could plausibly have many potential substrates on which to act. Thus, understanding the relative importance of these factors in regulatory evolution is critical for achieving a comprehensive view of expression divergence.

A common approach for detecting, and estimating the magnitudes of, *cis* and *trans* effects, is to measure allele-specific expression (ASE) in hybrids and their parents. ASE can be used to classify genes into regulatory types based on the presence and directionality of *cis* and *trans* components (McManus et al. 2010). It has been broadly applied to both intraspecific and interspecific hybrids to study the genetics of expression variation within and between species (reviewed in Gaur et al. 2013; Signor and Nuzhdin 2018). The comparison of within- and between-species regulatory genetics informs our understanding of evolutionary mechanisms because the concordance or discordance of phenomena on these two timescales can narrow the range of evolutionary explanations for the variation. A major conclusion from accumulated ASE research is that intraspecific gene expression evolution in animals is mediated predominantly through *trans* effects, while interspecific evolution proceeds predominantly via *cis* effects (reviewed in Signor and Nuzhdin 2018; Hill, Vande Zande, and Wittkopp 2021). A plausible interpretation of this pattern invokes variation in mutational opportunity and modes of selection. In this world view, *trans* regulation has a broader mutational target, but *trans*-effect mutations tend to have greater deleterious pleiotropic effects (P. J. Wittkopp 2005; Gruber et al. 2012; Lemos et al. 2008). Thus, such variants are common as low frequency polymorphisms, generating population level *trans* effects, but rarely fix. Alternatively, cis-acting variants may be less pleiotropic and therefore, more likely to fix. However, this pattern is not always observed. For example, Sánchez-Ramírez et al. (2021) found *cis*-regulatory divergence was more frequent than *trans-*regulatory divergence in male, but not female *Caenorhabditis* species. Some studies of flies have found relatively more *trans* effects between species (McManus et al. 2010; Coolon et al. 2014). Thus, the literature is still unsettled.

There is no reason to expect that the genetics of regulatory variation will be homogeneous across tissues in multicellular organisms, as the cell and developmental biology, as well as the influence of mutation and selection on expression phenotypes may vary across cell types, tissues, and organs. Indeed, empirical evidence supports the view that tissues and cell types exhibit varying rates of expression divergence (Gu and Su 2007; Brawand et al. 2011; Romero, Ruvinsky, and Gilad 2012; Kryuchkova-Mostacci and Robinson-Rechavi 2015; Liang et al. 2018; J. Chen et al. 2019; Pal, Oliver, and Przytycka 2021). Intraspecific studies of mice (Babak et al. 2015; Andergassen et al. 2017; St Pierre et al. 2022), humans (Babak et al. 2015; Leung et al. 2015; Castel et al. 2020), birds (Wang, Uebbing, and Ellegren 2017), and Drosophila (Combs et al. 2018), have revealed tissue-specific variance in *cis*-effects; genes may exhibit ASE in some tissues but not others, and the total number and magnitude of *cis*-effects also varies across tissues. These studies did not identify *trans-*effects, however, limiting the insight we have into how regulatory evolution varies among tissues.

Despite the importance of granular investigations of regulatory divergence, relatively little literature addresses the genetics of interspecific expression evolution at the level of organ or tissue. Indeed, much of the influential Drosophila literature on this topic analyzes parental and hybrid expression in whole animals (Patricia J. Wittkopp, Haerum, and Clark 2004; Landry et al. 2005; Patricia J. Wittkopp, Haerum, and Clark 2008; McManus et al. 2010; Wei, Clark, and Barbash 2014; Coolon et al. 2014) or heads (Graze et al. 2009). While this literature has generated important results and patterns, the possibility remains of henceforth undiscovered genetically and evolutionary important regulatory heterogeneity across organs in Drosophila (Ranz et al. 2023). Additional studies used ASE to investigate interspecific regulatory divergence in Drosophila testis (Haerty and Singh 2006; Lu et al. 2010; Llopart 2012; Brill et al. 2016; Banho et al. 2021); while this work has shed light on the regulatory basis of hybrid male sterility, conclusions from studies of this organ may not apply to gene regulatory evolution more broadly. Few studies of interspecific ASE in animals have investigated single somatic tissues and characterized both *cis* and *trans* regulatory divergence (Goncalves et al. 2012; Davidson and Balakrishnan 2016).

Our goal here is to contribute to the literature on the genetics of interspecific regulatory divergence using the accessory gland (AG) of Drosophila as a model. Seminal fluid proteins (Sfps) are secreted by the accessory glands, ejaculatory duct, and ejaculatory bulb and transferred to females along with sperm during mating and are essential for fertilization, similar to the seminal fluid of the mammalian prostate (reviewed in Poiani 2006; Wilson et al. 2017). The genus Drosophila has a polyandrous mating system featuring competition between males for matings, and between ejaculates for access to eggs (Boorman and Parker 1976; Imhof et al. 1998; Clark, Begun, and Prout 1999). Sfps of Drosophila and many other insects induce a range of physiological and behavioral changes in females comprising the post-mating response (PMR; reviewed in Ravi Ram and Wolfner 2007; Avila et al. 2011; Sirot et al. 2014; Wigby et al. 2020), including increased egg laying, facilitation of sperm storage, immune system responses, increased feeding rates, increased activity level and decreased sleep, and decreased receptivity to remating. PMR phenotypes evolve in response to sperm competition and sexual conflict (Hollis et al. 2019). Sfps play a key role in mediating sperm competition; genetic variation in Sfp loci is linked to competitiveness (Clark et al. 1995; Chapman et al. 2000; Fiumera, Dumont, and Clark 2005), and males respond to perceived level of competition through differential allocation of Sfps to the ejaculate (Sirot, Wolfner, and Wigby 2011; Hopkins et al. 2019).

Sfp protein sequences evolve at an especially rapid rate, often under the influence of recurrent directional selection (Tsaur, Ting, and Wu 1998; Aguadé 1999; Begun et al. 2000; Holloway and Begun 2004; Begun et al. 2006-3; Schully and Hellberg 2006; Wong et al. 2008; Majane, Cridland, and Begun 2022), perhaps partially due to sexual conflict (Swanson and Vacquier 2002; Haerty et al. 2007). Reduced selective constraint may also contribute to their rapid divergence (Dapper and Wade 2020; Patlar et al. 2021). Sfp genes have rapid rates of turnover (Wagstaff and Begun 2005; Mueller et al. 2005), exhibiting gene gain and loss even among closely related species (Begun and Lindfors 2005).

While there is a long history of work on sequence evolution and turnover in Sfps (reviewed in Hurtado et al. 2022), less is known about gene expression evolution among Sfps or in the accessory gland more broadly. RNAi knockdowns demonstrate that PMR phenotypes are sensitive to expression level of many Sfps (Ravi Ram and Wolfner 2007; Patlar and Civetta 2022), suggesting that Sfp expression is a plausible target of selection. Consistent with the observation that male-biased genes tend to have higher levels of interspecific expression divergence (Meiklejohn et al. 2003; Parisi et al. 2004; Ellegren and Parsch 2007; Brawand et al. 2011; Graveley et al. 2011; Assis, Zhou, and Bachtrog 2012; Whittle and Extavour 2019; Pal, Oliver, and Przytycka 2021), we recently reported rapid expression divergence as well as the evolution of novel genes and expression phenotype in the accessory gland (Cridland et al. 2020). However, Cridland et al. did not focus on the general properties of accessory gland transcriptome divergence, did not compare Sfp expression to expression divergence of other gene classes expressed in the accessory gland, and did not address the genetics of accessory gland expression divergence between species. To our knowledge, despite its prominent role in the Drosophila evolutionary genetics literature, there has been no formal analysis of regulatory evolution in the accessory gland.

While hybrid males derived from crosses between *D. melanogaster* and its sibling species, *D. simulans* are generally completely sterile or inviable (Sturtevant 1920), with severely atrophied or absent testes, a previous report noted that hybrids between male *D. melanogaster* and female *D. simulans* have morphologically normal accessory glands, which produce seminal fluid that can induce the PMR in females (Stumm-Zollinger and Chen 1988). In this study, we have several goals. First, we characterize expression divergence between *D. melanogaster* and *D. simulans* in this key somatic organ of male reproduction, including contrasts between evolutionary properties of Sfps and other genes expressed in the gland. Second, we use ASE analyses derived from measures of gene expression in accessory glands of pure species and their hybrids to estimate *cis* and *trans* expression effects in the accessory gland and ejaculatory duct; we quantify inheritance of expression phenotypes and characterize misexpressed genes that may be related to hybrid incompatibilities. Third, we investigate connections between regulatory evolution and divergence in upstream noncoding regions and protein sequence evolution. Finally, we integrate ATAC-Seq data to link changes in chromatin state with expression divergence. Given the relatively low cell type diversity in these tissues and the numerical dominance of the accessory gland main cell (Majane et al. 2022), many of the inferences from these data are likely due to main cell expression phenotypes.

## METHODS

### RNA-Seq

We performed RNA-Seq on each of three samples: *D. melanogaster* (Raleigh 517, Mackay et al. 2012), *D. simulans* (*w^501^*), and a *D. melanogaster* X *D. simulans* interspecific hybrid, with three biological replicates per sample. We raised all Drosophila stocks on cornmeal-molasses-agar medium at 25°C and 60% relative humidity, on a 12:12 light/dark cycle. We crossed the parents to produce interspecific hybrids by pairing five *D. simulans* females with 25 *D. melanogaster* males; the excess of males helps overcome behavioral isolation between species. We collected virgin male adult F1 animals and kept them in groups of five males per vial. We aged flies for two days before dissection. On the day of the experiment we anesthetized flies with light CO_2_, dissected their accessory glands and anterior ejaculatory duct in cold 1X PBS, and collected the tissue in TRIzol (Thermo-Fisher 15596026) on ice. We confirmed that the hybrid organs appeared morphologically normal with seminal fluid production. The cell diversity of the dissected tissue is low, with roughly 15% of cells deriving from the ejaculatory duct and 85% from the accessory gland, which is composed primarily of a single cell-type, the main cell (Majane et al. 2022). After we collected tissue from 20 males we flash-froze the TRIzol tubes containing tissue in liquid nitrogen and stored the material at −80C. We extracted RNA using the standard TRIzol protocol followed by DNAse digestion (Invitrogen AM1907) and clean-up with AMPure beads (Beckman-Coulter A63881). Novogene performed RNA-Seq library preparation and Illumina sequencing (150bp paired-end).

### Assigning species-of-origin

To determine the species-of-origin for each allele in the hybrid, we used an alignment-based approach relying on differences in the number of mismatches between the reads and each reference. We aligned each hybrid sample to each of two references, *D. melanogaster* (custom reference based on Flybase release 6.04 with SNPs included for strain RAL 517), and *D. simulans* (Princeton University, release 3.0), using HiSat2 and requiring a MAPQ score ≥ 30. We sorted reads using custom perl, bash and R scripts (github.com/alexmajane/hybridASE) into groups that mapped either to one reference uniquely or mapped to both references. For read pairs where at least one mate aligned uniquely, we assigned both reads to that species. For the remaining reads, we analyzed the number of nucleotide mismatches algorithmically to assign species-of-origin. Reads that aligned to one species with at least six fewer mismatches were assigned to the species with fewer mismatches. We also subjected our *D. melanogaster* and *D. simulans* samples to the same workflow, to A) account for artifactual effects of the procedure on expression analysis, B) establish a ground-truth false-positive rate for species-assignment, and C) identify problematic gene regions with high rates of erroneous species-assignment. To address the latter point, we calculated the fold change of counts (see below) per gene with and without inclusion of misassigned parental reads. If the absolute value of log_2_(fold change) was greater than 0.025 in either species, we removed that gene from our downstream analyses, a total of 382 genes.

### Quantification of gene expression

After we assigned hybrid reads to species-of-origin, we quantified gene expression, using reads with a confident species assignment, with Salmon (Patro et al. 2017). We used Salmon’s alignment-free approach because it accounts for differences in transcript length and GC content across samples, which in our case vary among orthologs across species due to evolution and/or gene annotation. Following quantification we used tximport (Soneson, Love, and Robinson 2015) to estimate counts per gene. We limited our analysis to *D. melanogaster*-*D. simulans* 1-to-1 orthologs based on the FlyBase annotation (02/2020 release) with additional manually curated Sfp orthologs (Majane, Cridland, and Begun 2022).

### Differential expression

We used estimated counts from tximport as the basis of all downstream analyses. We performed independent analyses of autosomal and X-linked genes because of their different inheritance in the hybrid. We used DESeq2 (Love, Huber, and Anders 2014) to normalize count data with the median-of-ratios method (Anders and Huber 2010), identified DE genes using Wald tests, and estimated moderated log-fold changes with the *apeglm* model (Zhu, Ibrahim, and Love 2019).

### Regulatory and inheritance classifications

We refer to counts from *D. melanogaster* as P*_mel_*, *D. simulans* as P*_sim_*, and allele-specific hybrid counts as F1*_mel_* and F1*_sim_*. We calculated total F1 expression (F1_total_) as F1_mel_ + F1_sim_. We classified genes into regulatory and inheritance categories using the algorithm following McManus et al. (2010). For this purpose we define DE as a significant Wald test (Bonferonni adjusted p < 0.05), and make comparisons between A) parental expression: P*_mel_* and P*_sim_* (**P**), B) ASE within the hybrid: F1*_mel_* and F1*_sim_* (**H**), and C) between parent expression and expression of parental-specific alleles in the hybrid: P*_mel_* and F1*_mel_* or P*_sim_* and F1*_sim_* (**T**). We define regulatory classes as follows:

A. conserved: no significant **P**, **H**, or **T**.
B. *cis*: significant **P** and **H**, no significant **T**.
C. *trans*: significant **P** and **T**, no significant **H**.
D. *cis* + *trans*: significant **P**, **H**, and **T**, same directionality between the parental contrast and hybrid ASE. *cis* and *trans* effects favor expression of the same allele.
E. *cis* by *trans*: significant **P**, **H**, and **T**, opposite directionality between the parental contrast and hybrid ASE. *cis* and *trans* effects favor expression of different alleles.
F. compensatory: significant **H** and **T**, but no significant **P**. *cis* and *trans* effects complement one another such that there has been no evolved expression difference among species.
G. ambiguous: all other patterns of expression without classical interpretations, such as no **P** or **H**, but significant **T** (evidence of *trans* effects that appear only in the hybrid).

We classified genes by inheritance comparing overall hybrid expression (F1_total_) to parental expression. Note that throughout, the term *dominance* is used in the phenotypic sense; we do not make inferences about the genetic basis of inheritance of expression phenotypes in the hybrid. We define the following inheritance classes:

A. conserved: no DE between F1_total_ and P*_mel_* or P*_sim_*.
B. additive: DE between F1_total_ and both P*_mel_* and P*_sim_*. F1_total_ has an intermediate expression level.
C. *mel* dominant: no DE between F1_total_ and P*_mel_*. DE between F1_total_ and P*_sim_*.
D. *sim* dominant: no DE between F1_total_ and P*_sim_*. DE between F1_total_ and P*_mel_*.
E. overdominant: DE between F1_total_ and both P*_mel_* and P*_sim_*. F1_total_ is expressed at a higher level than both parents.
F. underdominant: DE between F1_total_ and both P*_mel_* and P*_sim_*. F1_total_ is expressed at a lower level than both parents.

### SFPs and AG-biased genes

We defined sets of seminal fluid proteins (Sfps) and accessory gland (AG)-biased genes to investigate patterns of DE, regulation, and inheritance in these gene classes. We refer to Wigby et al. (2020) for annotation of Sfps; 208 Sfps were expressed in our dataset. To annotate AG-biased genes we obtained expression data from FlyAtlas2 (Leader et al. 2018), and calculated the index of tissue specificity, τ (Yanai et al. 2005), for all genes. We defined AG-biased genes as those that are more highly expressed in the accessory gland than all other tissues, and have τ ≥ 0.8. There are 378 AG-biased genes expressed in our dataset, 238 of which are non-Sfps, which we used for further analysis.

### GO analysis

We performed gene ontology (GO) enrichment analyses with Enrichr (Kuleshov et al. 2016). We defined background gene sets as all genes expressed in the data (log_2_(counts) ≥ 1). We used the Bioconductor *D. melanogaster* annotation (Carlson 2021) and queried terms from all three sub-ontologies (Biological Process, Molecular Function, and Cellular Component).

### Upstream sequence analysis

We obtained sequences spanning 1000 bp, 750 bp, and 500 bp upstream of each *D. melanogaster* TSS (Flybase annotation 6.41), and removed overlapping coding sequence. We then used BLAST (gap open penalty: 2; gap extension penalty: 1) to identify orthologous sequence in the *D. simulans* genome (Princeton University, release 3.0). We discarded sequences with more than one BLAST hit or overlapping alignments. Next we aligned *D. melanogaster* and *D. simulans* sequences using MUSCLE (Edgar 2004). We estimated the Kimura 2-parameter nucleotide substitution rate (Kimura 1980) for each upstream region using EMBOSS distmat (Rice, Longden, and Bleasby 2000). We also trimmed the 500 bp sequences to shorter lengths of 100 bp, 200 bp, and 300 bp upstream of the TSS and repeated estimation of substitution rate.

### Protein sequence evolutionary analysis

We analyzed sequence divergence by estimating synonymous (dS) and nonsynonymous (dN) substitution rates. We obtained the longest open reading frame per gene from FlyBase annotations (*D. melanogaster* 6.41, *D. simulans* 2.02), translated nucleotide sequences with EMBOSS transeq (Rice, Longden, and Bleasby 2000), and aligned amino acid sequences with MUSCLE (Edgar 2004). We back-translated to codon alignments with gaps removed using PAL2NAL (Suyama, Torrents, and Bork 2006). For each gene we estimated dN and dS using Goldman and Yang’s maximum likelihood codon-based substitution model (codeml; Goldman and Yang 1994). We performed this analysis with PAML (Yang 1997) implemented in BioPython (Cock et al. 2009).

We additionally analyzed adaptive protein evolution in *D. melanogaster* using available population genomics data (Fraïsse, Puixeu Sala, and Vicoso 2019) with pre-computed McDonald-Krietman tests (McDonald and Kreitman 1991), as in our previous work (Majane, Cridland, and Begun 2022- see supplement for detail). We used the summary statistic *α*, which estimates for each gene the proportion of amino acid substitutions due to positive selection. A positive value is consistent with directional selection; larger values suggest a greater proportion of adaptive substitutions.

### Chromatin state integration

We analyzed chromatin state in *D. melanogaster* and *D. simulans* with ATAC-Seq data from Blair et al. (unpublished), who inferred ATAC-Seq peaks called from three replicates each in the same strains that we used. We used reciprocal BLAST with gap open penalty = 2 and gap extension penalty = 1 on peak sequences between species and verified orthology of 1-to-1 best hits using synteny (nearest upstream and downstream annotated exons). We defined <u>conserved</u> peaks as those with a single reciprocal hit in each species and shared synteny. We inferred synteny from the nearest upstream and downstream exons. We defined <u>orphan</u> peaks as those with no BLAST hits. We additionally BLASTed orphan peak sequences to the reciprocal species’ genome, and filtered out peaks with A) no hits to the genome, B) a hit within 100 bp of any annotated peak, C) multiple hits. It is important to compare truly orthologous regions when we quantify peak accessibility by counting reads. We defined orthologous regions of orphan peaks in reciprocal species as the span of each BLAST hit. We re-annotated conserved peaks by reciprocal BLAST of peaks to each other species’ genome, and extended the boundaries of each peak to the span of BLAST hits intersecting the original annotation.

To quantify chromatin accessibility in each peak, we counted the number of aligned ATAC-Seq reads intersecting each peak with HTSeq (Anders, Pyl, and Huber 2015). We analyzed count data using DESeq2 similarly to RNA-Seq analysis. We then annotated peaks with the nearest TSS. For each gene, we used our Salmon quantification to select the transcript with highest expression in our sample, and chose its annotated TSS. We then selected 1-to-1 peak-to-TSS pairs with the closest or overlapping peak per TSS, removing any duplicate matches from further analysis. We also filtered the data to include only pairs where each orthologous peak (or region in the case of orphans) matched the same gene in both species.

## RESULTS

### Alignment and identification of allele-specific reads

We performed RNA-Seq on *D. melanogaster*, *D. simulans*, and an interspecific hybrid (P*_mel_*, P*_sim_*, and F1). We obtained between 25.4 and 30.8M reads per RNA-Seq sample. We aligned each sample to each of two references, *D. melanogaster* and *D. simulans.* Each species aligns at a rate of 94-97% to the matching reference (Supplemental Table 1). A slightly better alignment rate for P*_sim_* is expected, since the *D. simulans* strain we used in our experiment matches the reference strain. Alternatively, our *D. melanogaster* strain matches the updated reference sequence for SNPs but not for the small number of indels in the genic regions used here.

For the reads from hybrids a higher alignment rate to *D. simulans* is expected given that the hybrid inherits a *D. simulans* X chromosome. Reads from the F1 aligned to the *D. melanogaster* reference at a rate of 61-62% and to the *D. simulans* reference at a rate of 69-71%. Among F1 reads, ∼20% aligned uniquely to *D. melanogaster*, 33-34% aligned uniquely to *D. simulans*, and 46-47% aligned to both references (Supplemental Table 2). We used a mismatch-based approach to compare alignments and assign non-uniquely aligned reads to each species (Supplemental Table 3). We were able to assign ∼25% of non-uniquely aligned reads (∼12% of total aligned reads) to *D. melanogaster*, and 35-36% of non-uniquely aligned reads (16-17% of total aligned reads) to *D. simulans*, leaving 18-19% of total aligned reads of indeterminable origin and removed from our analysis.

We passed parental pure species reads through the same analysis pipeline used for the F1, to 1) account for artifactual effects of the procedure on expression analysis, 2) establish a ground-truth false-positive rate for species-assignment, and 3) identify problematic gene regions with high rates of erroneous species-assignment. ∼1.2% of P*_mel_* reads uniquely aligned to the *D. simulans* genome, and 0.6-0.7% of P*_sim_* reads uniquely aligned to the *D. melanogaster* genome (Supplemental Table 2). Among non-uniquely aligning parental reads, our assignment algorithm assigned 0.16-0.17% of P*_mel_* reads to *D. simulans*, and 0.031-0.035% of P*_sim_* reads to *D. melanogaster* (Supplemental Table 3). We used incorrectly assigned parental reads to identify gene regions with elevated levels of misassignment. Most misassigned reads do not overlap genes (Supplemental Table 4). We identified 382 genes where misassigned reads significantly impacted estimates of gene expression (Supplemental Data); we removed these from downstream analysis. We used Salmon (Patro et al. 2017) to quantify gene expression on reads that passed our filtering and species-assignment. Salmon mapping rates are as follows: 89-90% of F1 reads of *D. melanogaster* origin (hereafter F1*_mel_*), 84-85% of F1 reads of *D. simulans* origin (hereafter F1*_sim_*), ∼94% of P*_mel_* reads, and 86-87% of P*_sim_* reads.

### Transcriptome-wide view

We converted quantified expression values from Salmon to estimated counts with tximport (Soneson, Love, and Robinson 2015) and used these counts as the basis of all downstream analyses. Given the different inheritance patterns of the autosomes and X chromosome in the hybrid, we analyze the corresponding genes separately. The hemizygous X is omitted from the ASE analysis. We refer to total F1 expression (sum of both alleles) as F1_total_, and ASE measures as F1*_mel_* and F1*_sim_*.

Principal component analysis of transcriptome-wide gene expression shows a high level of shared variance between replicates (Figure 1).

**Figure 1.**
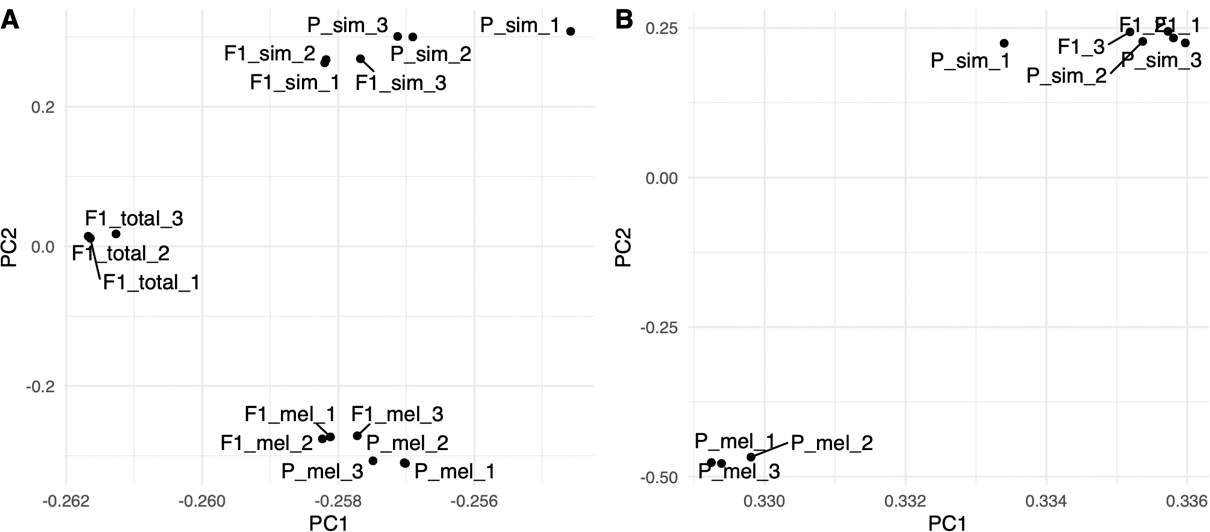
PCA of transcriptome-wide expression (log-transformed counts) showing the first two PCs. A) Autosomal-linked genes expression. Parental expression is overall similar to parent-specific alleles in the hybrid, with total hybrid expression clustering away from parental and allele-specific expression. B) X-linked gene expression. *D. simulans* expression is similar to hybrid expression.

Among autosomal genes, P*_mel_* groups with F1*_mel_* and P*_sim_* groups with F1*_sim_*, while F1_total_ groups away from these two clusters. PC1 appears to explain differences between F1 samples and P*_mel/sim_*, with allele-specific samples lying between F1_total_ and P*_mel/sim_*, though much closer to P*_mel/sim_*. PC2 appears to explain expression differences between *D. melanogaster* and *D. simulans*, with F1_total_ lying roughly midway between the species. Among X-linked genes, all of which derive from the *D. simulans* parent, both PC1 and PC2 appear to explain differences between *D. melanogaster* and *D. simulans* allele-derived expression, and X-linked F1 expression is tightly grouped with P*_sim_* expression. We did not find appreciable differences in distributions of expression variance between samples or allele-specific expression (Supplemental Figure 1), aside from a very modest elevation in standard deviation among X-linked genes relative to autosomal genes.

We characterize transcriptome-wide divergence using the correlations of average gene expression, which are high, as expected for two closely related species. Among autosomal genes, P*_mel_* and P*_sim_* have a Pearson correlation coefficient *r* = 0.934 (Figure 2A). F1_total_ expression is more similar to each parent; expression profiles between F1_total_ and P*_mel_* have an *r* = 0.965 (Figure 2B), while F1_total_ and P*_mel_* have an *r* = 0.967 (Figure 2C). Correlations between allele-specific expression and the same-species parent are strongest: F1*_mel_* and P*_mel_* have an *r* = 0.982 (Figure 2D); F1*_sim_* and P*_sim_* have an *r* = 0.980 (Figure 2E). Allele-specific expression within hybrids is somewhat more correlated than parental expression profiles are to one another; F1*_mel_* and F1*_sim_* have an *r* = 0.945 (Figure 2F). Among X-linked genes, P*_mel_* and P*_sim_* are similarly correlated as with autosomal genes (*r* = 0.930, Figure 2G). X-linked gene expression in hybrids is overall very similar to *D. simulans*: F1_total_ and P*_sim_* have an *r* = 0.983 (Figure 2H). Comparing F1_total_ and P*_mel_* is very similar to the parental X-linked expression contrast with an *r* = 0.929 (Figure 2I). Overall, the data strongly support the existence of widespread additivity for autosomal genes, and *D. simulans*-like expression on the hybrid X chromosome, suggesting strong influence of *cis* effects. The strong similarity in expression between the hybrid and the parents, and between hybrid ASE and parents, suggests that the accessory glands of this hybrid are not subject to particularly widespread misexpression.

**Figure 2.**
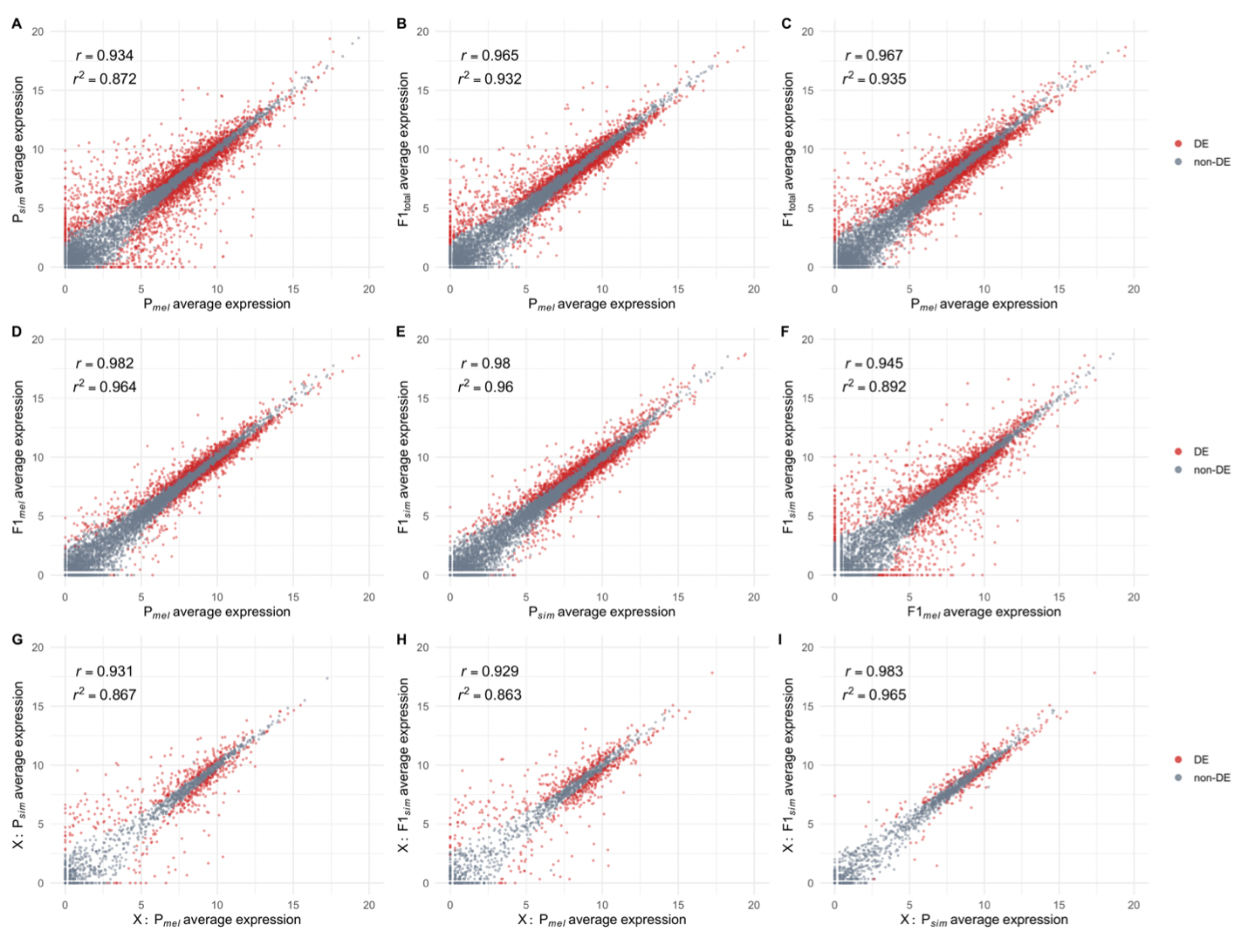
Corellations of average gene expression, *r* is the Pearson correlation coefficient, *r^2^* is the coefficient of determination in a simple linear model. (A-C) Comparisons of parental expression and total hybrid expression. (D-F) Comparisons of ASE. (G-l) Comparisons of X-linked gene expression

### Differential gene expression

Here we define differential expression as genes with a significant difference in normalized counts (Wald test, adjusted p < 0.01) and an absolute value of moderated log_2_(fold change) > 1 (Table 1). We also include counts of DE genes without imposing a log_2_(fold change) cutoff in Table 1. While many more genes are DE without a cutoff, in general we find that the patterns among these two criteria are similar, and given that DE genes with a fold-change cutoff are potentially more biologically relevant, we discuss them further below.

**Table 1.** Differentially expressed genes. DE is defined as a significant Wald test (Bonferroni adjusted p < 0.01), and a log_2_(fold change) value greater than 1. The rightmost three columns additionally show the number and fraction of significant DE genes without an imposed log_2_(fold change) cutoff.

| gene group | contrast | DE:<br>$\log_2(\text{FC}) > 1$ | non-DE:<br>$\log_2(\text{FC}) > 1$ | fraction DE:<br>$\log_2(\text{FC}) > 1$ | DE | non-DE | fraction DE |
| --- | --- | --- | --- | --- | --- | --- | --- |
| autosomes | $P_{mel} P_{sim}$ | 1608 | 7615 | 0.174 | 3005 | 6218 | 0.326 |
| autosomes | $F1_{total} P_{mel}$ | 883 | 8340 | 0.096 | 2290 | 6933 | 0.248 |
| autosomes | $F1_{total} P_{sim}$ | 956 | 8267 | 0.104 | 2459 | 6764 | 0.267 |
| autosomes | $F1_{mel} P_{mel}$ | 430 | 8793 | 0.047 | 1585 | 7638 | 0.172 |
| autosomes | $F1_{sim} P_{sim}$ | 561 | 8662 | 0.061 | 1790 | 7433 | 0.194 |
| autosomes | $F1_{mel} F1_{sim}$ | 1103 | 8120 | 0.120 | 2132 | 7091 | 0.231 |
| X | $P_{mel} P_{sim}$ | 284 | 1415 | 0.167 | 564 | 1135 | 0.332 |
| X | $F1_{sim} P_{mel}$ | 303 | 1396 | 0.178 | 631 | 1068 | 0.371 |
| X | $F1_{sim} P_{mel}$ | 66 | 1633 | 0.039 | 310 | 1389 | 0.182 |

We find that 17% of 9,223 total expressed genes are DE between P*_mel_* and P*_sim_* (Table 1). The list of DE genes and the resulting GO enrichments are provided in the Supplemental Data. The two most significantly enriched terms were GO:0005887 and GO:0031226, integral and intrinsic component of plasma membrane, respectively. Among the significant terms with at least two-fold enrichment we find neurotransmitter:sodium symporter activity (GO:0005328), solute:sodium symporter activity (GO:0015370), solute:cation symporter activity (GO:0015294), extracellular ligand-gated ion channel activity (GO:0005230, and neurotransmitter receptor activity (GO:0030594). Among the genes associated with such functions are multiple serotonin and octpamine receptors, as well as *nAchRB3*. The DE serotonin receptors, *5-HT1A*, *5-HT1B*, ands *5-HT2A*, are all expressed at substantially higher levels in *D. simulans*, and while the possible significance for AG function is unclear, such variation could plausibly be related to reproductive behavioral divergence between the two species, such as copulation duration (Grant 1983; Welbergen, Scharloo, and Van Dijken 1987; Price et al. 2001). Several gustatory and odorant receptors were DE, as were several odorant binding protein genes. A number of DNA damage response genes were also differentially expressed. As expected, fewer genes are DE in comparison to the hybrid: 10% of expressed genes are DE between F1_total_ and P*_mel_*, and 10% between F1_total_ and P*_sim_*.

DE is less frequent between hybrid allele-specific expression and parents relative to total hybrid expression (Table 1): 5% of genes are DE between F1*_mel_* and P*_mel_*, and 6% of genes are DE between F1*_sim_* and P*_sim_*. 12% of genes are DE between F1*_mel_* and F1*_sim_* (allele-specific expression within the hybrid, indicative of *cis*-regulatory effects). Among expressed X-linked genes, 17% are DE between P*_mel_* and P*_sim_*,similar to autosomal genes. DE is more common between F1 and P*_mel_* with 18% of X-linked genes DE. Just 4% of X-linked genes are DE between F1 and P*_sim_*, suggesting that *trans*-regulatory differences associated with large shifts in X-linked gene expression are rare.

### DE among seminal fluid proteins (Sfps) and accessory gland-biased genes

Sfps are known to have very high rates of amino acid substitutions (e.g., Swanson et al. 2001; Haerty et al. 2007) as well as gene turnover between species (Wagstaff and Begun 2005, Mueller et al. 2005, Begun and Lindfors 2005, Hurtado et al. 2022). Given the observation that rates of protein evolution are often correlated with gene expression evolution (Makova and Li 2003; Nuzhdin et al. 2004; Lemos et al. 2005; Khaitovich et al. 2005; Jordan, Mariño-Ramírez, and Koonin 2005; Liao and Zhang 2006; Sartor et al. 2006; Hunt et al. 2013; Warnefors and Kaessmann 2013; Hodgins et al. 2016; Zhong, Lundberg, and Råberg 2021), we asked whether expressed Sfps (208 total) were more likely than non-Sfps to be DE between *D. melanogaster* and *D. simulans* (P*_mel_* vs P*_sim_*) and between hybrid alleles (ASE: F1*_mel_* vs F1*_sim_*). Indeed, 28% of Sfps are DE between *D. melanogaster* and *D. simulans*, compared to 17% of non-Sfps (G test, p < 0.001, Table 2), and 22% of Sfps are DE between hybrid alleles (ASE), compared to 12% of non-Sfps (G test, p < 0.001).

**Table 2.** Number of DE genes among gene classes. *AG-biased* and *non-biased* gene sets exclude Sfps. DE is defined as a significant Wald test (Bonferroni adjusted p < 0.01), and a log_2_(fold change) value greater than 1. The rightmost three columns additionally show the number and fraction of significant DE genes without an imposed log_2_(fold change) cutoff.

| gene class | contrast | DE:<br>$\log_2(\text{FC}) > 1$ | non-DE:<br>$\log_2(\text{FC}) > 1$ | fraction DE:<br>$\log_2(\text{FC}) > 1$ | DE | non-DE | fraction DE |
| --- | --- | --- | --- | --- | --- | --- | --- |
| non-Sfp | $P_{mel} P_{sim}$ | 1549 | 7466 | 0.172 | 2883 | 6132 | 0.320 |
| Sfp | $P_{mel} P_{sim}$ | 59 | 149 | 0.284 | 122 | 86 | 0.587 |
| non-biased | $P_{mel} P_{sim}$ | 1461 | 7316 | 0.166 | 2733 | 6044 | 0.311 |
| AG-biased | $P_{mel} P_{sim}$ | 88 | 150 | 0.370 | 150 | 88 | 0.630 |
| non-Sfp | $F1_{mel} F1_{sim}$ | 1057 | 7958 | 0.117 | 2025 | 6990 | 0.225 |
| Sfp | $F1_{mel} F1_{sim}$ | 46 | 162 | 0.221 | 107 | 101 | 0.514 |
| non-biased | $F1_{mel} F1_{sim}$ | 989 | 7788 | 0.113 | 1889 | 6888 | 0.215 |
| AG-biased | $F1_{mel} F1_{sim}$ | 68 | 170 | 0.286 | 136 | 102 | 0.571 |

Sfps are a highly expressed class of genes, however, (Supplemental Figure 2A-C), and so may be more likely to be DE because of greater statistical power to detect expression differences. The median log_2_(counts) of Sfps is 8.46, while the median of non-Sfps is 4.31. To account for the effect of expression level on the likelihood of DE, we used a multiple logistic regression with average expression and Sfp status as independent variables and DE as the dependent variable (DE ∼ log_2_(counts) + Sfp). Interestingly, highly expressed genes are not much more likely to have large effect-size DE (log_2_(fold change) > 1); average expression level has a weak relationship with DE between P*_mel_* and P*_sim_* (β = 0.04 ± 0.01, p < 0.001). Sfp status does predict large-effect size DE (β = 0.45 ± 0.16, p = 0.006). Average expression does not have a relationship with DE between F1*_mel_* and F1*_sim_* (β = 0.01 ± 0.01, p = 0.08), and Sfp status predicts DE (β = 0.53 ± 0.17, p = 0.002). We therefore conclude that Sfps are significantly enriched for large-effect size DE events compared to non-Sfps. If we consider genes to be DE without a log_2_(fold change) cutoff however, we observe a strikingly different result. In the parental contrast, average expression significantly predicts DE between P*_mel_* and P*_sim_* (β = 0.22 ± 0.01, p < 0.001), however Sfp status does not predict DE (β = −0.04 ± 0.16, p = 0.792). Considering ASE, average expression significantly predicts DE between F1*_mel_* and F1*_sim_* (β = 0.21 ± 0.007, p < 0.001), but Sfp status does not predict DE (β = 0.09 ± 0.16, p = 0.554). Therefore we conclude that in our experiment, Sfps are not any more likely to be DE without a cutoff than non-Sfps when accounting for expression level.

Sfps have at least two obvious attributes; they are transferred to females during mating and they tend to exhibit AG-biased expression. To investigate whether the enrichment of Sfps amongst DE genes is more likely related to their status as Sfps or to their AG-biased expression, we also characterized DE in AG-biased genes. There are 238 genes that are AG-biased (τ > 0.8) but are not Sfps; 17.2% of non-biased genes are DE between P*_mel_* - P*_sim_*, and 11.7% are DE between hybrid alleles. The proportion of AG-biased genes exhibiting DE is dramatically greater; 37% are DE between P*_mel_* - P*_sim_*, and 28.6% are DE between hybrid alleles (Table 2). AG-biased genes are more highly expressed than non-AG-biased genes, but not as highly expressed as Sfps (Supplemental Figure 2D-F).

To ask whether AG-biased genes are more likely to be DE, we used a multiple logistic regression on non-Sfps, with average expression and AG-bias as independent variables, and DE as the dependent variable (DE ∼ log_2_(counts) + AG-bias). In the parental contrast, average expression only very weakly predicts DE between P*_mel_* and P*_sim_* (β = 0.03 ± 0.01, p < 0.001), while AG-bias very strongly predicts DE (β = 0.95 ± 0.14, p < 0.001). For ASE, average expression does not predict DE between F1*_mel_* and F1*_sim_* (β = 0.01 ± 0.01, p = 0.40), and AG-bias strongly predicts DE (β = 0.96 ± 0.15, p < 0.001). It is therefore apparent that AG-biased genes are much more likely to be DE than more broadly expressed genes in the accessory gland. Thus, there are two conclusions. First, from the expression divergence perspective, Sfps are not obviously different from other AG-biased genes, thereby providing no direct support for the idea that high Sfp expression divergence is a direct result of their function in the female reproductive tract (though our observations do not rule out this possibility). Second, while the role of directional selection in large interspecific expression differences cannot be inferred with these data, the enrichment pattern is consistent with the idea that expression phenotypes are more likely to be the result of selection on gland-related phenotypes rather than as a pleiotropic effect of selection on expression in other organs.

### Gene regulatory divergence classification

We characterized *cis-* and *trans*-regulatory effects for each autosomal gene by comparing ASE in hybrids to expression in each parent species. 2764 genes have evidence of *cis* effects (30% of expressed genes), 3338 have evidence of *trans* effects (36%), and 1601 have evidence of both *cis* and *trans* effects (17%). While there are more genes with significant *trans* effects (G test, p < 0.001), the median *cis* effect is significantly larger (log_2_(fold change) = 0.92) than the median *trans* effect (log_2_(fold change) = 0.64, Wilcoxon rank sum test, p < 0.001).

We further classified the regulatory basis of each gene following McManus et al. (2010) (Figure 3, Supplemental Tables 5, 6). The biggest category (4062 genes, or 44%) is <u>conserved,</u> with no significant *cis* or *trans* effects. Roughly equal numbers of genes exhibit pure *cis* or pure *trans* regulation; 933 = 10.1%, and 912 = 9.9%, respectively. Genes with both *cis* and *trans* effects are classified into three groups. The largest group by far is *<u>cis</u>* <u>+ *trans*</u> regulation, where *cis* and *trans* effects have the same directionality (eg. P*_mel_* > P*_sim_* and F1*_mel_* > F1*_sim_*), which contains 1116 (12.1%) genes. Only 104 (1.1%) genes have *<u>cis</u>* <u>by *trans*</u> regulation—*cis* and *trans* effects with opposite directionality (eg. P*_mel_* > P*_sim_* and F1*_mel_* < F1*_sim_*). Finally, 381 genes (4.1%) exhibit <u>compensatory</u> regulation, such that there is no DE between P*_mel_* and P*_sim_* despite evidence of *cis* and *trans* regulatory evolution. There are 1715 genes (18.6%) that cannot be classified into any of the above categories, and are labeled <u>ambiguous</u>.

**Figure 3.**
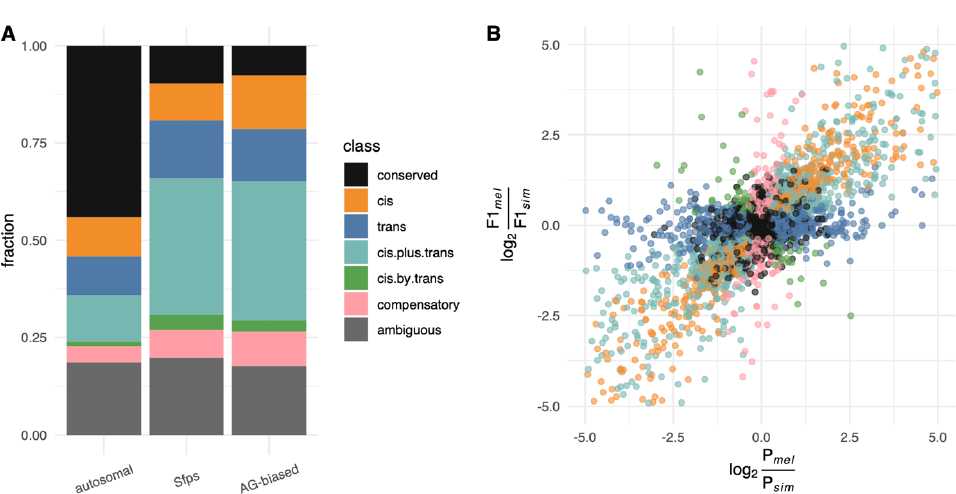
Regulatory classification by *cis* and *trans* mechanisms. A) Fraction of genes classified into each regulatory type, with Sfps and AG-biased genes shown separately. B) log_2_(fold change) of ASE and parental expression are shown with regulatory types highlighted. Ambiguous genes are removed and scale limited for clarity. For full data visualization see Supplemental Figure 3.

Sfps and AG-biased genes are much less likely to be conserved (Figure 3A, Supplemental Table 5), which is expected given their higher rates of expression divergence. Removing conserved and ambiguous classifications from consideration more clearly reveals differences in how Sfps and accessory gland-biased genes are regulated relative to all autosomal genes (Supplemental Table 6); both have smaller proportions of *cis*-regulation. There are about half as many *cis*-regulated genes among Sfps and a third fewer among accessory gland-biased genes. *cis* + *trans* regulation is particularly common among Sfps and accessory gland-biased genes. Rates of pure *trans*, *cis* by *trans*, and compensatory regulation are roughly equal among gene sets.

### Inheritance classification, misexpression, gain- and loss-of-function phenotypes

We characterized patterns of inheritance of expression phenotypes for autosomal genes by comparing F1_total_ to each parent (Figure 4, Supplemental Tables 7, 8). <u>Conserved</u> genes exhibit no DE in any comparison, comprising 4886 genes (53% of all expressed genes). 731 genes (7.9%) are <u>additive</u>, where P*_mel_* and P*_sim_* are DE and F1_total_ has an intermediate expression phenotype. Genes with parental divergence and with F1_total_ expression levels that were not DE relative to either parent are classified as either *<u>mel</u>* <u>dominant</u> or *<u>sim</u>* <u>dominant</u>. Rates are similar: 1403 (15.2%) are *mel* dominant and 1208 (13.1%) are *sim* dominant. Genes that are overexpressed in F1_total_ relative to both parents are <u>overdominant</u>, and underexpressed genes <u>underdominant</u>. There are 448 (4.9%) overdominant and 547 (5.9%) underdominant genes.

**Figure 4.**
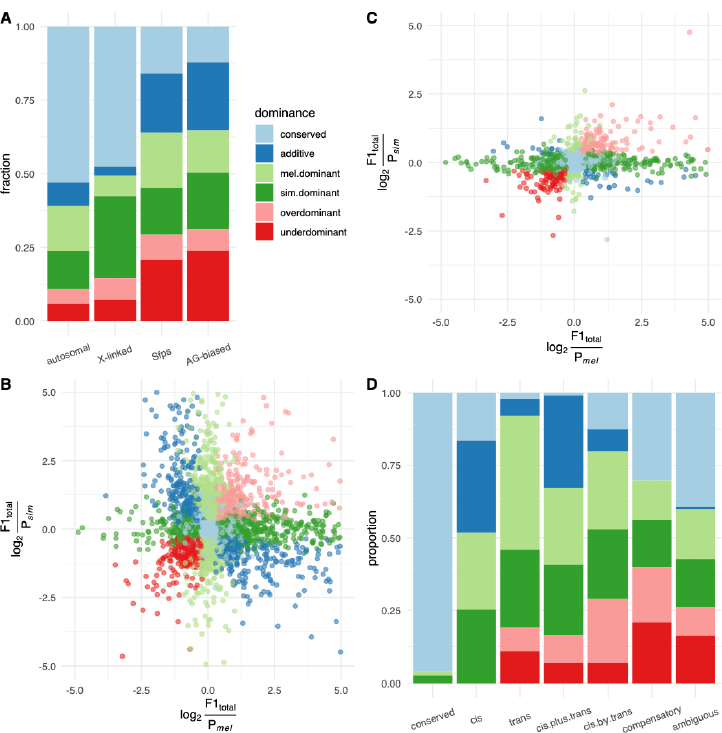
Inheritance classification gene expression phenotypes in hybrid offspring. A) Fraction of inheritance types for autosomal, X-linked, Sfps, and AG-biased genes are each shown independently. B) log_2_(fold change) of hybrid expression relative to each parent for autosomal genes. C) X-linked genes. Scale is limited in B-C for clarity. For full visualization, see Supplemental Figure 3. D) Inheritance types among each of the regulatory classifications identified through *cis* and *trans* mechanisms.

We also classified X-linked genes according to phenotypic inheritance patterns (Figure 4A,C, Supplemental Tables 7, 8). Compared to autosomal genes, X-linked have a similar percentage of conserved genes. As expected given the hemizygous *sim* X chromosome and lack of *cis* effects in the hybrid, there is little additivity (3%) or *mel* dominant inheritance (7%); X-linked genes have a strong excess of *sim*-dominant phenotypes (28%). X-linked genes are also more likely to be underdominant or overdominant compared to autosomal genes: 14.5% of X-linked genes are misexpressed compared to 10.8% of autosomal genes (G-test, p < 0.001), consistent with the faster-X hypothesis (Vicoso and Charlesworth 2006).

As with regulatory classes, Sfps and accessory gland-biased genes are much less likely to be conserved than all genes. Looking at the distributions of non-conserved classes (Supplemental Table 8), it is clear that both gene classes are more likely to be additive than *mel* or *sim* dominant. Sfps and accessory gland-biased genes additionally have significantly higher levels of underdominance than overdominance, a departure from trends among all autosomal genes.

Beyond misexpression, we also classified genes that are not DE between parents and which have a gain-of-function (GOF, overexpression in hybrids) or loss-of-function (LOF, underexpression in hybrids) expression phenotype (Supplemental Data). There are 58 genes with significant GOF (12% of all overexpressed genes), and 40 with significant LOF (10% of all underexpressed genes)—representing relatively rare events. Further, restricting GOF to cases with insignificant expression in parents (log_2_(counts) < 1 in P*_mel_* and P*_sim_*, log_2_(counts) > 1 in F1_total_) leaves only five instances, including *prolyl-4-hydroxylase-⍺ MP* and four uncharacterized genes (Supplemental Table 9). There are two cases of LOF with insignificant hybrid expression (log_2_(counts) > 1 in P*_mel_* and P*_sim_*, log_2_(counts) < 1 in F1_total_): *β-Tubulin at 85D* and *Pendulin* (Supplemental Table 10).

Next we examined the relationship between regulatory and inheritance classes (Figure 4D, Supplemental Table 11). We expect that genes with stronger *cis* components would be more likely to have an additive inheritance pattern, on the basis of the relative contributions of each species’ allele to total hybrid expression (Lemos et al. 2008; McManus et al. 2010). We found that genes with strong *cis* regulatory components had the highest levels of additivity: 31.8% of *cis* and 31.9% of *cis* + *trans* regulated genes had additive inheritance, compared to just 5.9% of *trans* regulated genes. We expected that genes with antagonistic *cis* and *trans* components would be more likely to lead to misexpression in hybrids, highlighting potential incompatibilities between species. Indeed, we find that *cis*-by-*trans* and compensatory gene classes are more likely than others to be associated with underdominant/overdominant inheritance. *Cis*-by-*trans* regulated genes have an excess of overdominance (22%) relative to underdominance (7%), but this may be attributable to the small sample size (n = 104 genes). Finally, we observe a strong trend towards *trans* regulated genes being inherited in a *mel*-dominant fashion (46% of *trans* regulated genes, compared to just 27% being *sim*-dominant; *mel*-dominant genes have a significantly greater proportion of *trans* regulation: G-test, p < 0.001).

### GO enrichment analysis

To investigate possible biological correlates of regulatory and inheritance classes we investigated the associated GO enrichments (Figure 5, Supplemental Figure 4, Supplemental Data). To increase the sample of X-linked genes, we used genes overexpressed or underexpressed in the hybrid relative to *D. simulans*, rather than genes strictly classified as overdominant or underdominant. Purely *cis*-regulated, *cis*-by-*trans*, and conserved genes are generally associated with larger p values and/or weakly enriched GO terms. Among our more significant results are 73 terms associated with translation in purely *trans*-regulated genes, driven by several ribosomal subunit and eukaryotic elongation factor genes (Supplemental Figure 4A, Supplemental Data). Translation-related genes are also significantly enriched among *mel*-dominant, and particularly for overdominant inheritance. Among the 40 translation proteins that are *trans* and *mel*-dominant, 32 are more highly expressed in *D. melanogaster* (chi-square test, p = 0.005).

**Figure 5.**
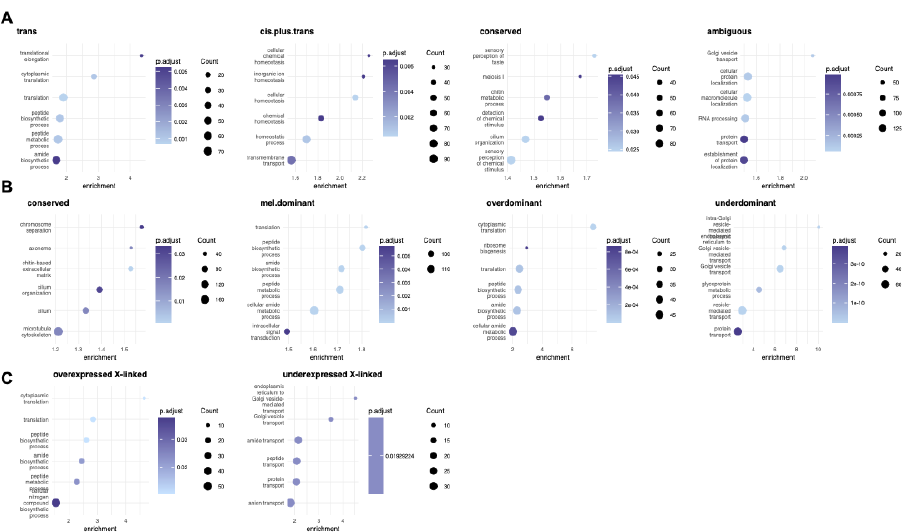
GO terms associated with regulatory and inheritance types. Enrichmentis the ratio of terms in the test gene set compared to the background gene set. A) Regulatory types; B) inheritance types in autosomes; C) over- or underexpressed X-linked genes. Terms associated with *cis* and *cis* by *trans* genes are weakly significant and not shown here (see Supplemental Data).

The remaining 33 genes exhibiting other modes of inheritance show no bias towards either parent (chi-square test, p = 0.82). There are 61 overdominant translation-related genes on the autosomes. This gene set only partially overlaps with the *trans*-regulated gene set—18 of overdominant translation-related genes are *trans*-regulated, but eight are *cis* + *trans*, six are *cis* by *trans*, 12 are compensatory, and 17 are ambiguous. Translation-related GO terms are also enriched in overexpressed genes on the X chromosome (Supplemental Figure 4B,C, Supplemental Data). On the X chromosome, there are an additional 52 overexpressed genes associated with translation. Taken together, the data suggest that translation-related genes are especially likely to be both *trans*-regulated and overdominant, but that overdominance may be the result of diverse regulatory mechanisms.

Underdominant inheritance / underexpression is strongly associated with golgi / endoplasmic reticulum vesicle transport GO terms on both the autosomes (137 genes, Supplemental Figure 4B) and X chromosome (36 genes, Supplemental Figure 4C). Of 137 underdominant transport-related genes on the autosomes, 86 have an ambiguous regulatory classification, while 20 are *trans*, 18 are *cis* + *trans*, and 13 are compensatory. Of the ambiguous terms, all are non-DE between parents, and also non-DE between hybrid alleles. Therefore there is no evidence of *cis* effects in these genes. Underdominance is indicative of *trans* factors, however these effects have not led to divergence between *D. melanogaster* and *D. simulans*, suggesting *trans* effects that occur specifically in the hybrid.

### Upstream sequence divergence

To investigate the possible sequence basis of expression variation we characterized divergence in the regions upstream of AG-expressed genes. We analyzed distributions of substitution rate for various upstream sequence lengths (Supplemental Figure 5); the 300 bp region captured the highest overall levels of divergence, so we chose this set for further analysis. We observe significant variation in upstream sequence evolution among regulatory and inheritance classes (Kruskal tests, p < 0.01). Among regulatory classes, conserved genes have the lowest rate of upstream sequence divergence with a median of 0.070 substitutions/bp (Figure 6A). All other classes except *cis* by *trans* have significantly greater divergence rates (Wilcoxon rank sum tests, p < 0.01). Ambiguous and compensatory genes have the greatest rates at 0.088 substitutions/bp. Among inheritance classes, additive and conserved genes are very similar; median = 0.074 per bp and 0.073 per bp, respectively (Wilcoxon rank sum test p > 0.05). Alternatively, underdominant genes have much higher rates of upstream sequence divergence with a median = 0.104/bp (pairwise Wilcoxon rank sum tests, p < 0.001 vs all other classes). Given the enrichment of underdominant genes for golgi/protein transport-related GO terms, we asked whether those genes were confounded with the elevated level of upstream sequence divergence. Of 451 underdominant genes with upstream sequence information, 109 are associated with golgi / protein transport-related GO terms. If we remove these from the analysis, underdominant genes still have significantly greater upstream sequence divergence than all other classes (median = 0.101/bp, pairwise Wilcoxon rank sum tests, p < 0.001). Genes with *mel* or *sim* dominance exhibit intermediate levels of upstream sequence divergence (medians = 0.086 and 0.080/bp, respectively). Since *cis*-regulatory evolution could proceed through mutations in promoter regions, we asked whether the magnitude of ASE or parental divergence is correlated with upstream sequence divergence, however, we observed no relationship (Supplemental Figure 6). Upstream sequence divergence in Sfps or AG-biased genes does not differ significantly from non-Sfps / non-AG-biased genes (Wilcoxon rank sum tests, p = 0.16, p = 0.26, respectively). Overall, then, the most obvious correlate of upstream sequence divergence is hybrid misexpression.

**Figure 6.**
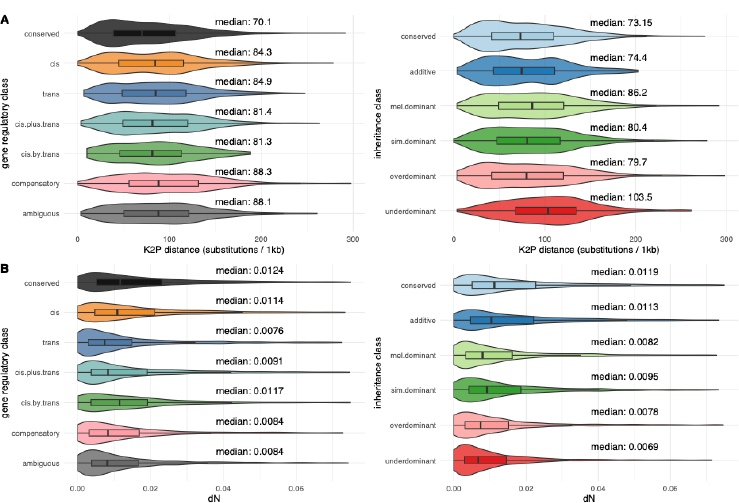
A) Distributions of Kimura-2-parameter estimated substitution rates among regulatory and inheritance classes. B) Distributions of nonsynonomous substitution rate (dN) among regulatory and inheritance classes. K2P distance and dN vary significantly across (Kruskal tests, p < 0.001). Alongside the median, significant differences by pairwise Wilcoxon rank sum tests (Holm-Bonferroni adjusted p < 0.05) are indicated by different numbers across gene sets.

### Protein sequence evolution

Previous studies have found positive correlations between gene expression divergence and protein sequence evolution (Makova and Li 2003; Nuzhdin et al. 2004; Lemos et al. 2005; Khaitovich et al. 2005; Jordan, Mariño-Ramírez, and Koonin 2005; Liao and Zhang 2006; Sartor et al. 2006; Hunt et al. 2013; Warnefors and Kaessmann 2013; Hodgins et al. 2016; Zhong, Lundberg, and Råberg 2021). We investigated whether this was the case for AG-expressed genes. Additionally, we investigated whether rates of protein sequence evolution differ among regulatory and inheritance classes.

We observed no association between protein sequence evolution (dN) and expression divergence between parents or ASE (Supplemental Figure 7A,B). Additionally, we did not observe a strong difference between dN and r regulatory or inheritance classes (Figure 5B). A multivariate regression of dN by expression level and parental expression divergence suggests that genes with parental conservation have greater dN than DE genes, though effect sizes are very small (average expression: β = 1.3×10^−3^ ± 8.1×10^−5^, p < 0.001; parental expression DE: β = 1.7×10^−3^ ± 6.4×10^−4^, p = 0.007). Notably, we observe that genes with higher expression levels tend to have lower dN, consistent with the literature (Pál, Papp, and Hurst 2001; reviewed in Drummond et al. 2005).

Median dN is 3.6 times greater for Sfps than non-Sfps (Wilcoxon rank sum tests, p < 0.001). Among AG-biased non-Sfps, dN is modestly elevated, 1.3 times higher, but still significantly different from non-accessory gland-biased genes (Wilcoxon rank sum tests, p = 0.00101, p < 0.001)(Supplemental Figure 7C,D). Increased protein divergence could be explained by directional selection (Tsaur, Ting, and Wu 1998; Aguadé 1999; Begun et al. 2000; Holloway and Begun 2004; Begun et al. 2006-3; Schully and Hellberg 2006; Wong et al. 2008; Majane, Cridland, and Begun 2022) or reduced constraint (Dapper and Wade 2020; Patlar et al. 2021). To seek evidence bearing on these alternatives we used McDonald-Kreitman tests and compared the summary statistic *α* (higher *α* suggests a greater overall proportion of adaptive amino acid substitutions) among gene classes. The resulting patterns are similar to—though weaker than—patterns in dN (Supplemental Figure 8A). In both regulatory and inheritance classes, conserved genes have significantly higher median *α* than some other types, but we do not observe significant differences among other classifications. As with dN, we observe significantly elevated *α* among Sfps (Supplemental Figure 8B): median Sfp *α* = 0.256; non-Sfp median *α* = −0.375 (Wilcoxon rank sum test, p < 0.001). Unlike dN, *α* does not differ significantly among AG-biased and non-biased genes (Supplemental Figure 8C, Wilcoxon rank sum test, p = 0.52). Thus, while Sfps and AG-biased non-Sfps tend to exhibit similar patterns of regulatory divergence, recurrent adaptive protein divergence appears to be more prevalent among Sfps.

### Chromatin state integration

We used ATAC-Seq data (Blair et al., unpublished) from *D. melanogaster* and *D. simulans* to investigate connections between chromatin accessibility and DE or ASE. We annotated ATAC-Seq peaks as <u>conserved</u>, *<u>mel</u>* <u>orphans</u>, or *<u>sim</u>* <u>orphans</u>. Conserved peaks are called in orthologous regions of both species, whereas orphan peaks are called only in one species.

In total we annotated 7,416 conserved, 2,370 orphan *sim*, and 1,680 orphan *mel* peaks. We made peak-to-gene associations annotating peaks to the closest / overlapping transcription start site (TSS), which left us with 2,898 conserved, 1,627 orphan *sim*, and 1,232 orphan *mel* peaks with 1-to-1 gene annotations. Among annotated conserved peaks, 88% overlap the TSS. Roughly 6% of non-overlapping peaks are upstream of the TSS, and 6% are downstream. The median width of conserved peaks is 644 bp in both *D. melanogaster* and *D. simulans*, while the mean is 759.5 bp in *melanogaster* and 758 bp in *simulans*. Orphan peaks are much less likely to overlap with the TSS: 16% of orphan *mel* and 19% of orphan *sim* have overlap. Orphan peaks are also more likely to be upstream than downstream. In *D. melanogaster*, 57% of orphan peaks are upstream while 26% are downstream; in *D. simulans*, 50% are upstream and 30% are downstream. Orphan peaks are also smaller than conserved peaks: the median width of *mel* orphans is 399 bp, while the median width of *sim* orphans is 255 bp. PCA of log_2_(counts) shows that replicates cluster together by species (Supplemental Figure 9A-C), though we note that clustering is not as strong as RNA-Seq data, which is expected due to the background and variance inherent of ATAC-Seq data.

Relative accessibility among species in conserved peaks is normally distributed (Supplemental Figure 10A; 25% percentile log_2_(*mel* / *sim*) = −0.205; 75% percentile = 0.212), suggesting there is no systematic directionality in chromatin accessibility between species. The distribution of log_2_(*mel* / *sim*) for orphan peaks is highly skewed towards each respective species (Supplemental Figure 10B,C; *mel* orphans: 25% percentile log_2_(*mel* / *sim*) = 0.48; 75% percentile = 1.50; *sim* orphans: 25% percentile log_2_(*mel* / *sim*) = −1.48; 75% percentile = −0.48, in line with the expectation that the species with a peak called will have higher accessibility. There are a small number of cases where the species without a peak has higher accessibility (2.3% of *mel* orphans, 4.9% of *sim* orphans). We removed from further consideration A) orphan peaks that are less accessible in the species with a peak present, and B) orphan peaks that are not DA, leaving 74% of *mel* and 72% of *sim* orphans (Supplemental Figure 10D,E).

There is a weak positive relationship between accessibility and expression (Figure 6A,B), suggesting that chromatin state and expression are indeed correlated. Given this relationship, we asked whether the presence of chromatin peaks was associated with the likelihood of DE in nearby genes. In both parental and ASE contrasts, DE genes are enriched for the presence of nearby orphan peaks relative to non-DE genes (Figure 6C,D, Supplemental Tables 12, 13). We used a multiple logistic regression with average expression and peak status as independent variables and DE as the dependent variable (DE ∼ log_2_(counts) + peak). In the parental contrast, presence of a conserved peak negatively predict DE (β = −0.17 ± 0.08, p = 0.046), and orphan peaks are strong predictors of DE (*mel* orphan: β = 0.78 ± 0.11, p < 0.001, *sim* orphan: β = 0.86 ± 0.09, p < 0.001). Similarly, conserved peaks are negatively correlated with DE in ASE (β = −0.30 ± 0.10, p = 0.002), and orphan peaks strongly predict DE (*mel* orphan: β = 0.85 ± 0.11, p < 0.001, *sim* orphan: β = 0.77 ± 0.10, p < 0.001). Average gene expression also predicts DE in both contrasts, but the regression coefficients are notably smaller in comparison with orphan peak presence (parental: β = 0.22 ± 0.01, p < 0.001; ASE: β = 0.21 ± 0.01, p < 0.001).

We further investigated correlates of gene expression divergence with presence or absence of nearby 1-to-1 peaks by comparing log_2_(fold changes) (Figure 6G). In the parental contrast (P*_mel_* - P*_sim_*), genes without a peak annotation have a narrower distribution of log_2_(P*_mel_* / P*_sim_*) relative to peaks with a conserved peak nearby, but both gene sets have medians near 0. The median absolute value of log_2_(P*_mel_* / P*_sim_*) in genes with a conserved peak is 0.37, significantly higher than genes with no peak, median = 0.29 (Wilcoxon rank sum test, p < 0.001). Genes with a *mel* orphan peak nearby are biased towards positive values of log_2_(P*_mel_* / P*_sim_*) with median = 0.252, and genes with a *sim* orphan peak nearby are biased towards negative values with median = −0.221. The median absolute values of log_2_(P*_mel_* / P*_sim_*) of genes near orphan peaks are significantly greater than genes near a conserved peak or no peak (median *mel* orphan = 0.61; *sim* orphan = 0.59; Kruskal test, p < 0.001, pairwise Wilcoxon rank sum tests, p < 0.001 in each case). Medians associated with *mel* peaks and *sim* peaks are not significantly different (Wilcoxon rank sum test, p = 0.30). We observe the same patterns in ASE among different classes of peaks, but the magnitude of expression differences is smaller compared to parental DE (Figure 7H).

**Figure 7.**
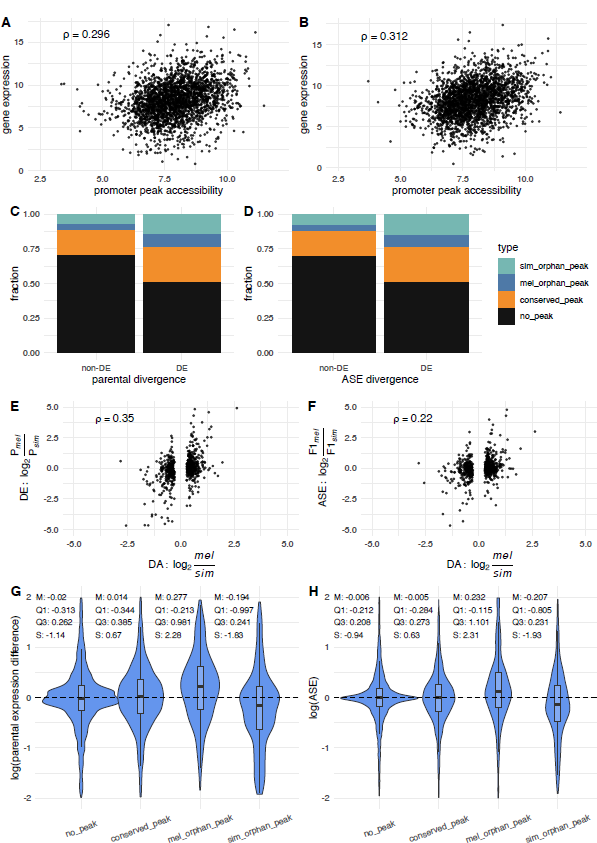
Interfacing promoter region accessibility estimated from ATAC-Seq data with gene expression. Gene expression and conserved peak accessibility have a positive relationship in (A) P*_mel_* and (B) P*_sim_*. Spearman’s rank coefficient ⍴ is displayed. DE genes are more likely to be associated with orphan peaks in both the (C) parental and (D) ASE contrast. (E) Parental expression divergence and (F) ASE (y-axis) for are plotted against accessibility differences for DA conserved chromatin peaks called by ATAC-Seq (x-axis). (G) Distributions of log2(fold change) of parental expression difference differ among peak types. M: median, Q1: 1^st^ quartile, Q3: 3^rd^ quartile, S: skewness. Genes associated with conserved peaks have a broader 1^st^-3^rd^ interquartile range and significantly larger median absolute value than genes without an annotated promoter peak. Orphan peaks are skewed towards greater expression values in the species with a peak. (H) Distributions of log2(fold change) of ASE show similar patterns to parental divergence.

Next, we asked whether the magnitude of expression differences between species or ASE was correlated with the magnitude of peak accessibility differences among DA peaks. We find a positive relationship between ranks of these measures (Figure 6E,F; conserved peaks, parental divergence: ⍴ = 0.35; ASE: ⍴ = 0.22). These correlations are, however, much weaker for orphan peaks (Supplemental Figure 11). In summary, it appears that the presence of accessible chromatin is correlated with differential expression of nearby genes, and that quantitative differences in conserved peak accessibility are correlated with concordant differences in gene expression. However, there is no strong quantitative relationship for orphan peaks.

To investigate the relationship between chromatin accessibility and *cis/trans* regulatory divergence we compared Spearman rank correlations between expression divergence and accessibility divergence among conserved, pure *cis*, pure *trans*, and *cis* + *trans* regulated genes (Supplemental Table 14). As expected, conserved genes exhibited no correlation of accessibility divergence with either parental expression (⍴ = −0.01) or ASE divergence (⍴ = −0.08). Genes with *trans*-regulatory components exhibit a moderately stronger correlation of accessibility divergence with parental expression divergence (⍴ = 0.52 for both *trans* and *cis* + *trans*) than pure *cis*-regulated genes (⍴ = 0.41). Genes with *trans*-regulatory components have a no correlation of accessibility divergence with ASE divergence (⍴ = 0.14), while *cis* (⍴ = 0.42) and *cis* + *trans* (⍴ = 0.44) genes have relatively strong correlations. It therefore appears that genes with *cis* and *trans* regulatory divergence have relatively stronger correlations between accessibility and parental expression divergence, and that only *cis*-regulated genes show correlations between accessibility divergence and ASE divergence.

Regulatory and inheritance classes are associated with different proportions of chromatin peak types (Figure 7, Supplemental Table 15). As expected, genes that are annotated as conserved in regulation or inheritance are much less likely to be associated with nearby peaks. The set of genes with purely *cis* regulation have a smaller proportion of conserved peaks than genes with significant *trans* factors. Genes with *cis* + *trans* regulation, and genes with additive inheritance, have the highest respective shares of orphan peaks. Underdominant genes have the highest share of conserved peaks among inheritance classes.

## DISCUSSION

Given that *D. melanogaster* and *D. simulans* are sister species that share a most recent common ancestor only 2-3 million years ago (Obbard et al. 2012), we expect strongly correlated phenotypes for most traits. In agreement with this expectation, autosomal gene expression profiles between *D. melanogaster* and *D. simulans* were very similar, as evidenced by transcriptome-wide expression correlations (*r* = 0.934). This correlation is somewhat stronger than that observed in our recent bulk RNA-Seq (Cridland et al. 2020) and single-cell RNA-Seq studies (Majane, Cridland, and Begun 2022) of the same tissues. Given the myriad technical differences between studies, it is difficult to interpret variation in the magnitudes of these correlations. Similarly, the lack of similar datasets for several organs in these two species, subjected to the same analysis pipeline, makes it difficult to draw strong conclusions about whether rates of transcriptome divergence in the AG are unusually large (Cridland et al. 2020).

Hybrid expression profiles are overall more similar to each parent than the parents are to each other and are roughly midpoint between parental expression in PCA. ASE profiles within the hybrid are more similar than parents are to one another, as expected, given that parental divergence is the result of *cis* and *trans* effects, while divergence between hybrid alleles is driven only by *cis* effects. Similarly, we observed a higher proportion of DE genes between parents than between parental alleles in the F1. As we see with transcriptomic correlations, large effect-size DE between hybrid alleles and parent-of-origin is rare (4% of genes), suggesting that *trans*-effects of large size are uncommon.

X-linked and autosomal gene expression profiles have very similar correlations between parents and also exhibit similar levels of DE. These results are unexpected since previous whole-animal transcriptome data shows a strong faster-X effect on expression divergence, even among non-male biased genes (Meisel, Malone, and Clark 2012; Kayserili et al. 2012). We observed a modest rate of DE between the hybrid and *D. simulans* X chromosomes, suggesting that while *trans* effects on the X are not very rare, the effect-sizes are generally small.

Since Sfps exhibit rapid protein divergence (Tsaur, Ting, and Wu 1998; Aguadé 1999; Begun et al. 2000; Holloway and Begun 2004; Begun et al. 2006-3; Schully and Hellberg 2006; Wong et al. 2008; Majane, Cridland, and Begun 2022) and genomic turnover (Wagstaff and Begun 2005; Mueller et al. 2005; Begun et al. 2006-3; Hurtado et al. 2022), we formally investigated their interspecific expression divergence. Sfps are much more likely to have large effect-size DE, consistent with previous work on major expression divergence in this tissue (Cridland et al. 2020). However, accessory gland-biased non-Sfps exhibit similar patterns. Thus, genes that make the largest contribution to the unique transcriptome of the accessory gland, regardless of whether they are Sfps, tend to exhibit greater expression divergence. Determining whether or not this property of the AG transcriptome is driven by directional selection is an important unsolved problem.

The median *cis* effect is 43% larger than the median *trans* effect, consistent with strong correlations of expression between hybrid alleles and their parents of origin. In contrast, McManus et al. (2010) observed significantly larger *trans* effects in whole female Drosophila hybrids. However, despite their smaller mean effect-size, *trans* effects are more common than *cis* effects. Some evidence suggests that *cis* effects contribute more to species expression divergence than trans-effects, perhaps as a consequence of their lower average pleiotropy (reviewed in Signor and Nuzhdin 2018; Hill, Vande Zande, and Wittkopp 2021; Ranz et al. 2023). Our results do not support this model, in line with results from some other studies (McManus et al. 2010; Coolon et al. 2014; Sánchez-Ramírez et al. 2021). Additional investigation of diverse somatic organs to understand the distribution of *cis* and *trans* effects across tissues, which could help answer a broader question of which factors influence the variation observed in relative levels of interspecific *cis* and *trans* divergence.

We classified genes into regulatory and inheritance classes as originally outlined in McManus et al. (2010). Equal proportions of genes have evolved through purely *cis* or trans *regulation*, with a larger proportion exhibiting evidence of both mechanisms, consistent with a complex basis of regulatory divergence. Opposing directionality of cis and trans evolution (*cis* by *trans* and compensatory classes) is rare in our study. Other studies in Drosophila found that opposing *cis* and *trans* effects were much more common (Ranz et al. 2004; Landry et al. 2005; Graze et al. 2009; McManus et al. 2010; Coolon et al. 2014). Studies of mouse liver (Goncalves et al. 2012) and testis (Mack, Campbell, and Nachman 2016) also have relatively high levels of opposing *cis* and *trans* effects. Fraser (2019) found that errors in estimation of ASE can lead to inflated estimates of *cis* by *trans* effects, however, suggesting that comparisons of these effects across studies may be complicated by technical issues. Sfps and accessory gland-biased genes appear to accumulate higher levels of cis + trans regulation than other AG-expressed autosomal genes, with purely cis or trans regulation being relatively rare. Additionally, Sfps and accessory gland-biased genes are more likely to exhibit additive inheritance than all other genes, as well as elevated misexpression (see below).

Misexpression in the hybrid AG is relatively rare, with just 10.8% of autosomal genes overdominant or underdominant. Further, just a handful of genes have complete GOF or LOF expression phenotypes. These results suggest that the accessory gland is not prone to widespread dysgenesis between these species, in contrast to results from Drosophila testis or female whole-animal data (Ranz et al. 2004; Haerty and Singh 2006; Moehring, Teeter, and Noor 2007; McManus et al. 2010; Coolon et al. 2014; Cartwright and Lott 2020; Ranz et al. 2023), Caenorhabditis data (Sánchez-Ramírez et al. 2021), mouse liver (Goncalves et al. 2012) and testis (Mack, Campbell, and Nachman 2016) data. Thus, our transcriptomic data are consistent with the observations of Stumm-Zollinger and Chen (1988), that hybrid accessory glands have relatively normal morphology, seminal fluid, and ability to induce the female PMR. Other studies have found limited levels of misexpression in Drosophila larva (Moehring, Teeter, and Noor 2007; Wei, Clark, and Barbash 2014), female heads (Graze et al. 2009), Hawai’ian Drosophila testes (Brill et al. 2016), and avian brains (Davidson and Balakrishnan 2016). Clearly, the level of hybrid dysgenesis in gene expression is highly variable among species and tissues; more work on tissue-specific ASE is needed to reveal the proximate and ultimate explanation for this variation.

Several Drosophila studies have reported that male-biased genes are prone to misexpression in hybrids, and are especially likely to be underdominant in whole animals or testis (Haerty and Singh 2006; Michalak and Noor 2003; Moehring, Teeter, and Noor 2007; McManus et al. 2010), a pattern observed in most, but not all Drosophila crosses (Banho et al. 2021). Underdominant male-biased genes are also linked to male sterility (Michalak and Noor 2004). We also found higher levels of misexpression among AG-biased genes, with a particular enrichment for overdominance. Autosomal genes have similar rates of overdominance and underdominance. Sfps have 2.7 times as many misexpressed genes compared to non-Sfps, with a ratio of underdominant to overdominant expression of 2.3; accessory gland-biased non-Sfps are have 2.9 times as many misexpressed genes, with a ratio of underdominant to overdominant expression of 3.4. These data suggest that the widely-reported observation of male biased underexpression is not limited just to the testis. Whether this pattern holds for male-biased genes across multiple somatic tissues or only those related to reproduction is an important question for future studies.

The faster-X hypothesis (Vicoso and Charlesworth 2006) predicts not only elevated expression divergence among X-linked genes, but also increased X-linked misexpression in hybrids. Faster-X divergence has been observed in flies (Meisel, Malone, and Clark 2012; Kayserili et al. 2012), but Drosophila hybrids actually have a lower rates of X misexpression in the testes (Lu et al. 2010; Llopart 2012) or no difference from autosomal misexpression in larvae (Wei, Clark, and Barbash 2014). In contrast, our data reveal no evidence of faster-X divergence, but do show faster-X misexpression, similar to patterns observed in mice (Good et al. 2010; Larson et al. 2016). Faster-X gene expression patterns may therefore vary among tissues in Drosophila, and future studies of tissue-specific hybrid gene expression are needed to determine the extent and biology underlying these potential differences.

Genes that are autosomal overdominant or X-linked overexpressed are both highly enriched for translation-related genes, including numerous elongation factors and ribosomal subunits. Overexpressed proteins in *D. melanogaster* - *D. simulans* hybrid embryos were enriched for “translation initiation” genes (Bamberger et al. 2018), and misexpressed genes in hybrid house mice testes were also significantly enriched for translation-related GO terms (Mack, Campbell, and Nachman 2016). Therefore, misexpression of translation-related genes could plausibly be related to hybrid incompatibilities across species and developmental stages. Notably, hybrid male sterility evolves very quickly and the regulation of Drosophila spermatogenesis occurs primarily at the translational level (Schäfer et al. 1995). Whether reduced hybrid male fertility and misexpression of translational machinery are functionally linked remains a speculative matter. Translation is also enriched among *trans*-regulated and *mel*-dominant genes in our data, but these gene sets are only partially overlapping with one another, suggesting that translation-related genes may be regulated and inherited via diverse mechanisms in the accessory gland, and that overdominant translation-related genes are not regulated through a unifying mechanism. Underdominant autosomal and underexpressed X-linked genes are highly enriched for genes related to protein transport, golgi, and endoplasmic reticulum. Notably, ambiguous regulated genes are also enriched for these GO terms, and this gene list substantially overlaps with underdominance. Ambiguous regulation may occur in many ways and is difficult to put into biological context. However, in this case, the vast majority of these genes are not DE between the parents but do have evidence of *trans*-effects leading to underdominance. This suggests that emergent properties of *trans* factors active specifically in hybrid cells leads to underexpression, potentially indicative of hybrid incompatibilities related to protein transport in the golgi and endoplasmic reticulum.

We analyzed levels of nucleotide divergence in regions upstream of the TSS, which could plausibly affect expression divergence. While we do not find a quantitative relationship between expression divergence and upstream sequence divergence, genes that are conserved in their regulation and inheritance tend to have a lower level of divergence. Compensatory, ambiguous, and particularly underdominant genes have elevated levels of upstream sequence divergence. This suggests that underdominance might be arising from incompatibilities between rapidly evolving *cis*-acting sequences and *trans* regulatory factors.

As expected, Sfps have much higher dN than non-Sfps. Sfps also have elevated median α, suggesting overall greater levels of adaptive substitutions in these genes. While accessory gland-biased non-Sfps also have elevated dN, they do not exhibit significant elevation in α. Thus, from a regulatory perspective, Sfps are not obviously different from other AG-biased genes. However, Sfps appear to experience much more recurrent adaptive protein divergence than other AG-biased genes. This supports the idea that proteins that interact directly with the external environment are more likely to evolve adaptively (Gillespie and Langley 1974). Somewhat unexpectedly, we found that among AG-expressed genes, those in the conserved regulatory category have the highest overall levels of dN, ω, and α. This suggests that for most genes expressed in the accessory gland, there is a modest decoupling between expression evolution in the gland and rapid protein evolution driven by selection.

We associated ATAC-Seq peaks with nearest TSS to identify chromatin tied to putative regulatory regions. We found that orphan peaks were highly biased in accessibility towards one species and observed a modest but significant correlation between gene expression and accessibility among conserved peaks, similar to results of some studies (Nair et al. 2021), but weaker than that observed in others (Starks et al. 2019). We observe weaker correlations with orphan peaks, which may be explained by the greater median distance of orphan peaks to the TSS. Additionally, the ATAC-Seq data we used here includes only accessory gland tissue, while the RNA-Seq data derive from the accessory gland and ejaculatory duct. We therefore expect that our observations relating gene expression and log_2_(fold changes) across groups to chromatin state in this study are conservative estimates of true relationships.

We further identified a weakly positive quantitative relationship between differential accessibility of conserved peaks and the log(fold-change) of DE, in contrast to some other studies that found stronger associations (Racioppi, Wiechecki, and Christiaen 2019; Gontarz et al. 2020; Nair et al. 2021; Sanghi et al. 2021). However, limiting the analysis to genes with evidence of *cis* or *trans* regulatory divergence, reveals a relatively strong correlation between species differences in peak accessibility divergence and gene expression divergence. Orphan peaks do not appear to have this quantitative relationship, but the presence of an orphan peak very strongly predicts both small- and large-effect size DE in nearby genes. Taken together, the data suggest that presence/absence of chromatin peaks (either by evolutionary gain or loss, which is impossible to determine with this data) likely contributes to gene expression differences between *D. melanogaster* and *D. simulans*—if a peak appears in one species, there is a better chance that the nearest gene will be DE, and more often than not in the direction of the species with the peak—but there is no evidence of a straightforward quantitative relationship, at least that we can detect with this dataset. To overcome some of the technical and biological variables that complicate this analysis, an allele-specific multi-omic gene expression and ATAC-Seq experiment on single cells (Cao et al. 2018; S. Chen, Lake, and Zhang 2019) would provide stronger insights into the relationships between chromatin state, expression, and regulation.

## ACKNOWLEDGEMENTS

This work was supported by National Institutes of Health grant NIGMS R35GM134930 to DJB and by a National Science Foundation Graduate Research Fellowship to ACM. The content is solely the responsibility of the authors and does not necessarily represent the official views of the National Institutes of Health.

## SUPPLEMENTAL TABLES

**Supplemental Table 1.**
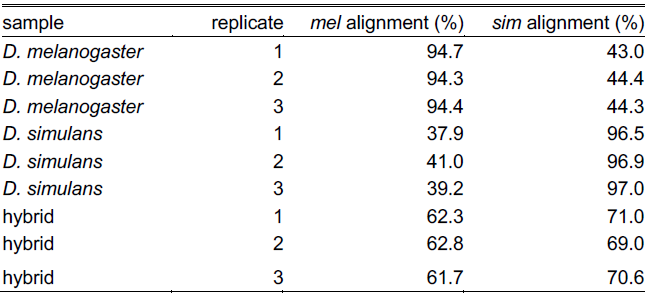
Overall alignment rates of reads to each reference genome: *D. melanogaster* (mel) and *D. simulans* (sim).

| sample | replicate | mel alignment (%) | sim alignment (%) |
| --- | --- | --- | --- |
| *D. melanogaster* | 1 | 94.7 | 43.0 |
| *D. melanogaster* | 2 | 94.3 | 44.4 |
| *D. melanogaster* | 3 | 94.4 | 44.3 |
| *D. simulans* | 1 | 37.9 | 96.5 |
| *D. simulans* | 2 | 41.0 | 96.9 |
| *D. simulans* | 3 | 39.2 | 97.0 |
| hybrid | 1 | 62.3 | 71.0 |
| hybrid | 2 | 62.8 | 69.0 |
| hybrid | 3 | 61.7 | 70.6 |

**Supplemental Table 2.** Uniquely mapping and shared reads among references. M = millions of reads.

| sample | replicate | mel unique | mel unique (%) | sim unique | sim unique (%) | shared | shared (%) |
| --- | --- | --- | --- | --- | --- | --- | --- |
| *D. melanogaster* | 1 | 11.93M | 45.4 | 0.32M | 1.20 | 14.1M | 53.4 |
| *D. melanogaster* | 2 | 11.51M | 43.6 | 0.33M | 1.23 | 14.6M | 55.1 |
| *D. melanogaster* | 3 | 12.9M | 44.1 | 0.36M | 1.22 | 16.0M | 54.7 |
| *D. simulans* | 1 | 0.16M | 0.62 | 13.1M | 52.3 | 11.8M | 47.1 |
| *D. simulans* | 2 | 0.17M | 0.69 | 11.8M | 50.9 | 12.4M | 48.4 |
| *D. simulans* | 3 | 0.17M | 0.63 | 13.8M | 50.6 | 13.3M | 48.7 |
| hybrid | 1 | 5.18M | 20.1 | 8.68M | 33.7 | 11.9M | 46.7 |
| hybrid | 2 | 5.56M | 20.5 | 9.02M | 33.3 | 12.5M | 46.1 |
| hybrid | 3 | 5.01M | 19.9 | 8.35M | 33.1 | 11.9M | 47.0 |

**Supplemental Table 3.** Species-of-origin assignments to reads that mapped to both genomes.

| sample | replicate | assigned mel (%) | assigned sim (%) | undetermined (%) |
| --- | --- | --- | --- | --- |
| *D. melanogaster* | 1 | 62.9 | 0.17 | 36.9 |
| *D. melanogaster* | 2 | 61.9 | 0.20 | 37.2 |
| *D. melanogaster* | 3 | 61.5 | 0.16 | 38.4 |
| *D. simulans* | 1 | 0.035 | 64.9 | 35.1 |
| *D. simulans* | 2 | 0.035 | 65.2 | 34.8 |
| *D. simulans* | 3 | 0.031 | 65.0 | 35.0 |
| hybrid | 1 | 24.7 | 34.8 | 40.4 |
| hybrid | 2 | 25.1 | 35.0 | 39.8 |
| hybrid | 3 | 25.6 | 35.6 | 38.7 |

**Supplemental Table 4.** Percentage of misassigned parental reads that do not overlap any gene features (corresponding to intergenic regions). Uniquely misassigned reads originate in the step of aligning reads to both genomes. Algorithmically misassigned genes aligned to both genomes but were misassigned when assigning species-of-origin.

| sample | replicate | uniquely misassigned<br>with no feature (%) | algorithmically misassigned with<br>no feature (%) |
| --- | --- | --- | --- |
| *D. melanogaster* | 1 | 73.7 | 39.5 |
| *D. melanogaster* | 2 | 72.0 | 41.1 |
| *D. melanogaster* | 3 | 73.9 | 41.7 |
| *D. simulans* | 1 | 53.6 | 45.8 |
| *D. simulans* | 2 | 54.1 | 45.4 |
| *D. simulans* | 3 | 53.1 | 44.6 |

**Supplemental Table 5.** Regulatory classifications among gene sets.

| regulatory class | *cis* | *trans* | *cis* + *trans* | *cis* by *trans* | compensatory | conserved | ambiguous |
| --- | --- | --- | --- | --- | --- | --- | --- |
| autosomal | 933 | 912 | 1116 | 104 | 381 | 4062 | 1715 |
| autosomal (%) | 10.12 | 9.89 | 12.1 | 1.13 | 4.13 | 44.04 | 18.59 |
| Sfps | 20 | 31 | 73 | 8 | 15 | 20 | 41 |
| Sfps (%) | 9.62 | 14.9 | 35.1 | 3.85 | 7.21 | 9.62 | 19.71 |
| AG-biased | 33 | 32 | 85 | 7 | 21 | 18 | 42 |
| AG-biased (%) | 13.87 | 13.45 | 35.71 | 2.94 | 8.82 | 7.56 | 17.65 |

**Supplemental Table 6.** Regulatory classifications (except conserved and ambiguous) among gene sets.

| regulatory class | *cis* | *trans* | *cis* + *trans* | *cis* by *trans* | compensatory |
| --- | --- | --- | --- | --- | --- |
| autosomal | 933 | 912 | 1116 | 104 | 381 |
| autosomal (%) | 27.07 | 26.47 | 32.39 | 3.02 | 11.06 |
| Sfps | 20 | 31 | 73 | 8 | 15 |
| Sfps (%) | 13.61 | 21.09 | 49.66 | 5.44 | 10.20 |
| AG-biased | 33 | 32 | 85 | 7 | 21 |
| AG-biased (%) | 18.54 | 17.98 | 47.75 | 3.93 | 11.80 |

**Supplemental Table 7.** Inheritance classifications among gene sets.

| inheritance class | additive | *mel* dominant | *sim* dominant | overdominant | underdominant | conserved |
| --- | --- | --- | --- | --- | --- | --- |
| autosomal | 731 | 1403 | 1208 | 448 | 547 | 4886 |
| autosomal (%) | 7.93 | 15.21 | 13.10 | 4.86 | 5.93 | 52.98 |
| X-linked | 51 | 119 | 473 | 125 | 121 | 810 |
| X-linked (%) | 3.00 | 7.00 | 27.84 | 7.36 | 7.12 | 47.68 |
| Sfps | 42 | 39 | 33 | 18 | 43 | 33 |
| Sfps (%) | 20.19 | 18.75 | 15.87 | 8.65 | 20.67 | 15.87 |
| AG-biased | 55 | 34 | 46 | 17 | 57 | 29 |
| AG-biased (%) | 23.11 | 14.29 | 19.33 | 7.14 | 23.95 | 12.18 |

**Supplemental Table 8.** Inheritance classifications (except conserved) among gene sets.

| inheritance class | additive | *mel* dominant | *sim* dominant | overdominant | underdominant |
| --- | --- | --- | --- | --- | --- |
| autosomal | 731 | 1403 | 1208 | 448 | 547 |
| autosomal (%) | 16.85 | 32.35 | 27.85 | 10.33 | 12.61 |
| X-linked | 51 | 119 | 473 | 125 | 121 |
| X-linked (%) | 5.74 | 13.39 | 53.21 | 14.06 | 13.61 |
| Sfps | 42 | 39 | 33 | 18 | 43 |
| Sfps (%) | 24 | 22.29 | 18.86 | 10.29 | 24.57 |
| AG-biased | 55 | 34 | 46 | 17 | 57 |
| AG-biased (%) | 26.32 | 16.27 | 22.01 | 8.13 | 27.27 |

**Supplemental Table 9.** Gain-of-function in hybrids with insignificant parental expression. Average expression is log_2_(normalized counts)

| gene | average expression in<br>*D. melanogaster* | average expression<br>in *D. simulans* | average expression in<br>hybrid |
| --- | --- | --- | --- |
| *prolyl-4-hydroxylase-α MP* | 0.000 | 0.000 | 2.681 |
| *CG9395* | 0.407 | 0.202 | 2.362 |
| *CG8500* | 0.705 | 0.706 | 2.324 |
| *CG30090* | 0.000 | 0.178 | 1.929 |
| *CG34316* | 0.795 | 0.380 | 2.501 |

**Supplemental Table 10.**
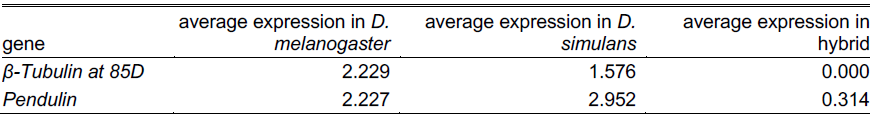
Loss-of-function with insignificant hybrid expression. Average expression is log_2_(normalized counts).

**Supplemental Table 11.**
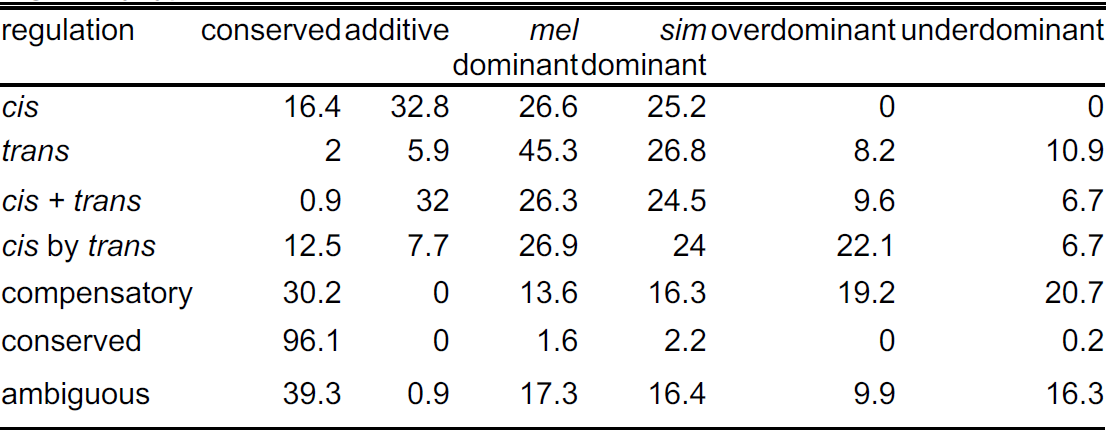
Percentage of inheritance classifications among regulatory types.

**Supplemental Table 12.**
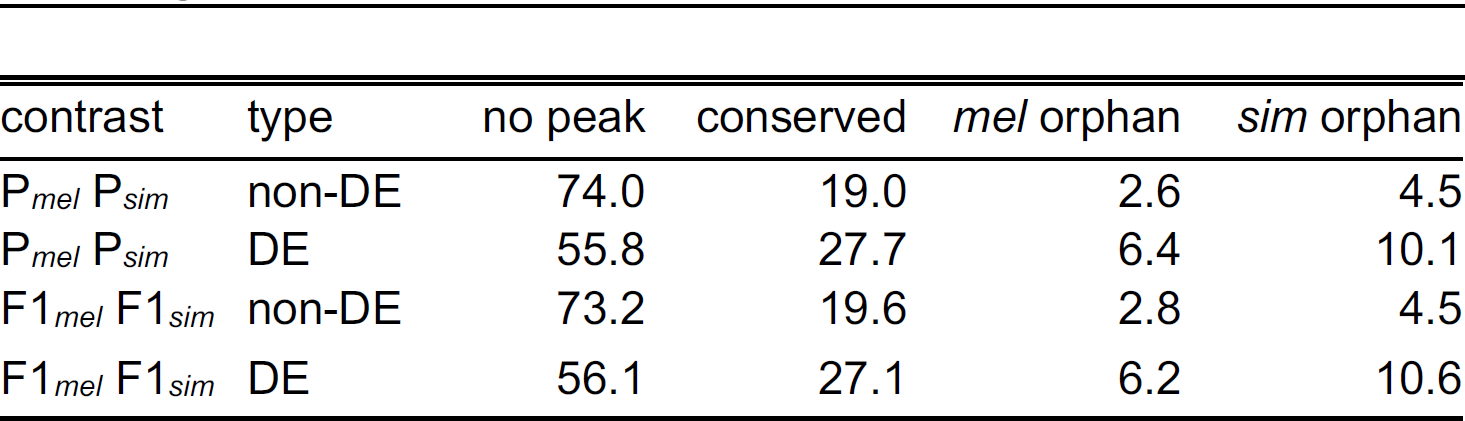
Percentage of chromatin peak types associated with DE genes.

**Supplemental Table 13.**
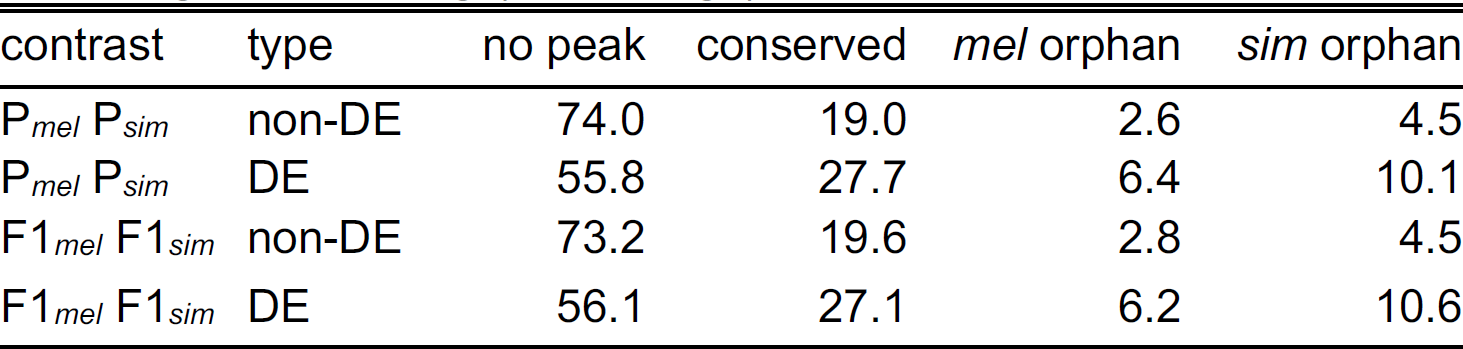
Percentage of chromatin peak types associated with DE genes, with log_2_(fold change) > 1.

**Supplemental Table 14.**
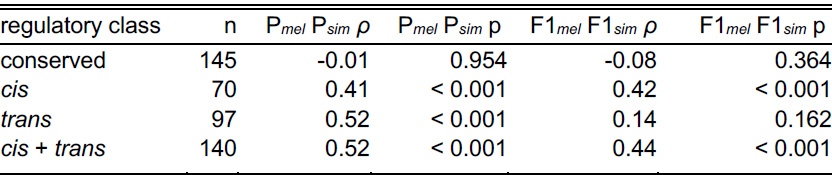
Correlations of expression divergence and accessibility divergence among regulatory classes, with differentially abundant, conserved ATAC-Seq peaks (as in Figure 6E,F). n is the sample size of genes in each comparison. P*_mel_* P*_sim_ ρ* is the Spearman rank coefficient of determination between parental expression divergence and chromatin accessibility divergence. P*_mel_* P*_sim_* p is the p value where the null hypothesis is that *ρ* = 0. Similarly, F1*_mel_* F1*_sim_ ρ* and p represent the Spearman rank coefficient of determination and p value for the correlation between ASE divergence and chromatin accessibility divergence.

**Supplemental Table 15.** ATAC-Seq peak classes among regulatory types.

| regulation | conserved peak | conserved peak (%) | *mel* orphan | *mel* orphan (%) | *sim* orphan | *sim* orphan (%) | no peak | no peak (%) |
| --- | --- | --- | --- | --- | --- | --- | --- | --- |
| *cis* | 209 | 22.4 | 83 | 8.9 | 99 | 10.6 | 542 | 58.1 |
| *trans* | 276 | 30.3 | 75 | 8.2 | 117 | 12.8 | 444 | 48.7 |
| *cis* + *trans* | 303 | 27.2 | 103 | 9.2 | 173 | 15.5 | 537 | 48.1 |
| *cis* by *trans* | 34 | 32.7 | 7 | 6.7 | 17 | 16.3 | 46 | 44.2 |
| compensatory | 111 | 29.1 | 27 | 7.1 | 57 | 15.0 | 186 | 48.8 |
| conserved | 527 | 13.0 | 111 | 2.7 | 182 | 4.5 | 3242 | 79.8 |
| ambiguous | 479 | 27.9 | 104 | 6.1 | 159 | 9.3 | 973 | 56.7 |

**Supplemental Figure 1.**
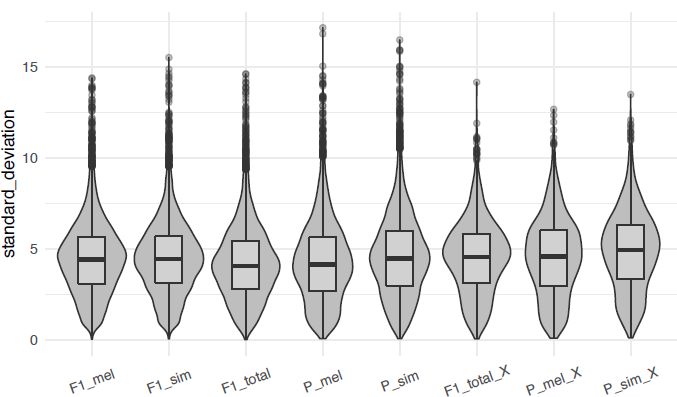
Distributions of observed standard devation of expression among replicates. The first five depict autosomal-linked genes and the last three depict X-linked genes.

**Supplemental Figure 2.**
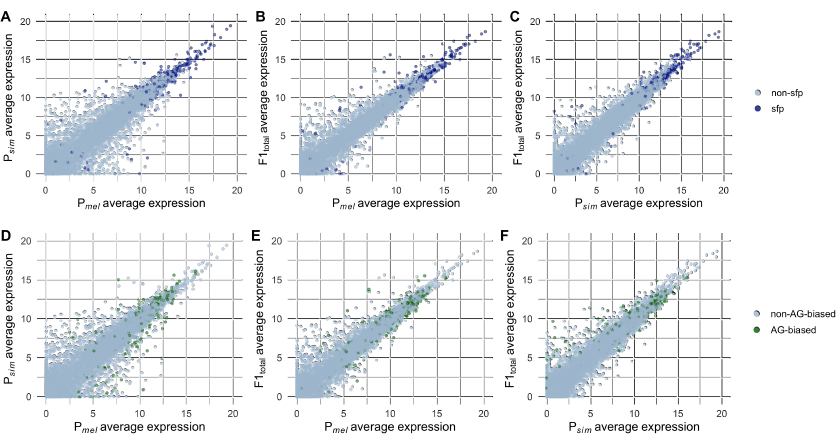
Corellations of average gene expression across the transcriptome, as in Figure 2, ith Sfps (A-C) and AG-biased genes (D-F) highlighted.

**Supplemental Figure 3.**
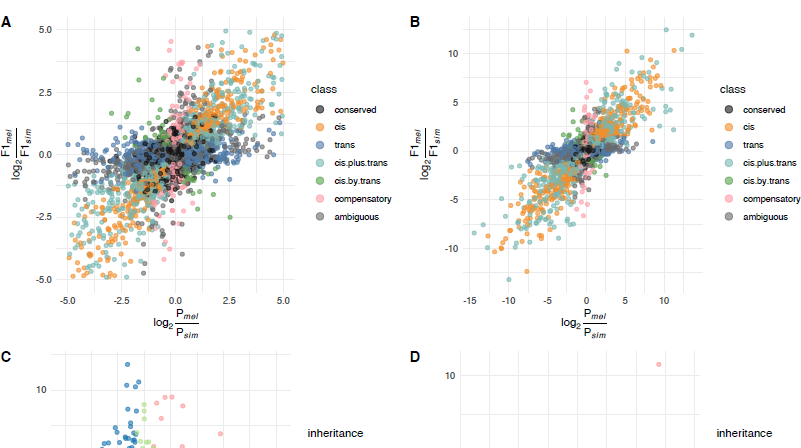
A) Regulatory classification as in Figure 3 with the addition of ambiguous genes. B) Full-scale log_2_(fold change) regulatory plot. C) Full-scale log_2_(fold ange) inheritance plot for autosomal-linked genes. D) Full-scale log_2_(fold change) heritance plot for X-linked genes.

**Supplemental Figure 4.**
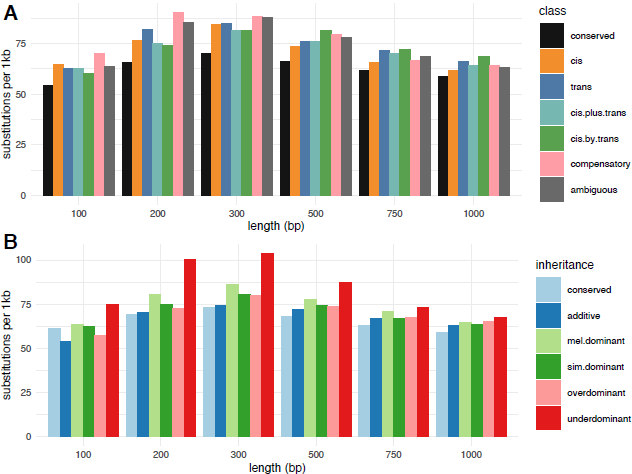
Kimura-2-parameter estimated substitution rates in upstream sequences between *D. melanogaster* and *D. simulans* at various lengths from the TSS. A) regulatory classes; B) inheritance classes.

**Supplemental Figure 5.**
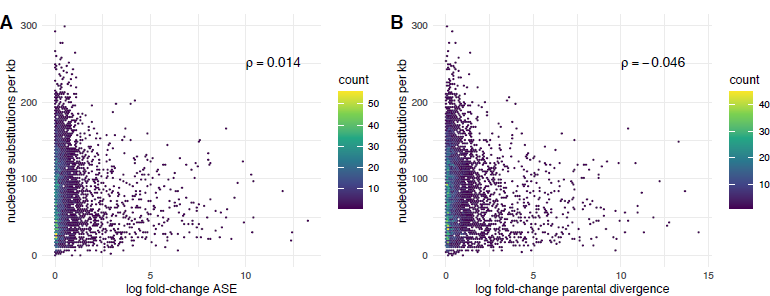
Kimura-2-parameter estimated substitution rates in upstream sequences between *D. melanogaster* and *D. simulans* plotted against A) allele-specific and B) parental expression divergence. Spearman’s rank correlation coefficient ⍴ is shown on each panel.

**Supplemental Figure 6.**
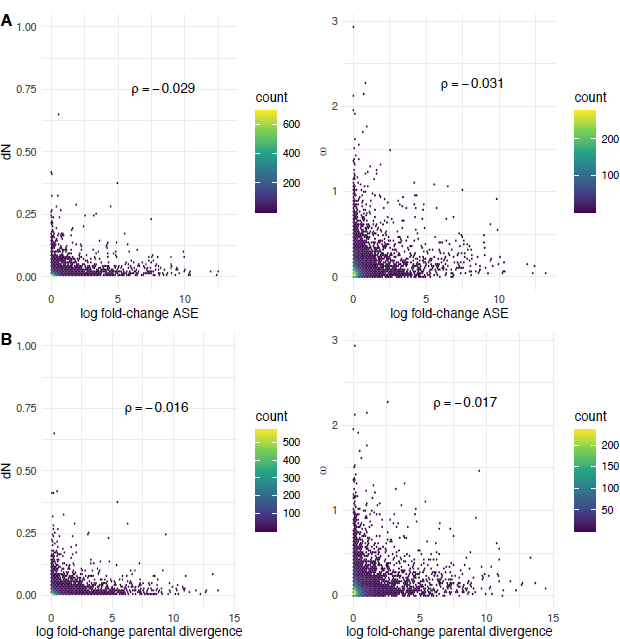
dN and ω between *D. melanogaster* and *D. simulans* plotted against A) allele-specific and B) parental expression divergence. Spearman’s rank correlation coefficient ⍴ is shown on each panel.

**Supplemental Figure 7.**
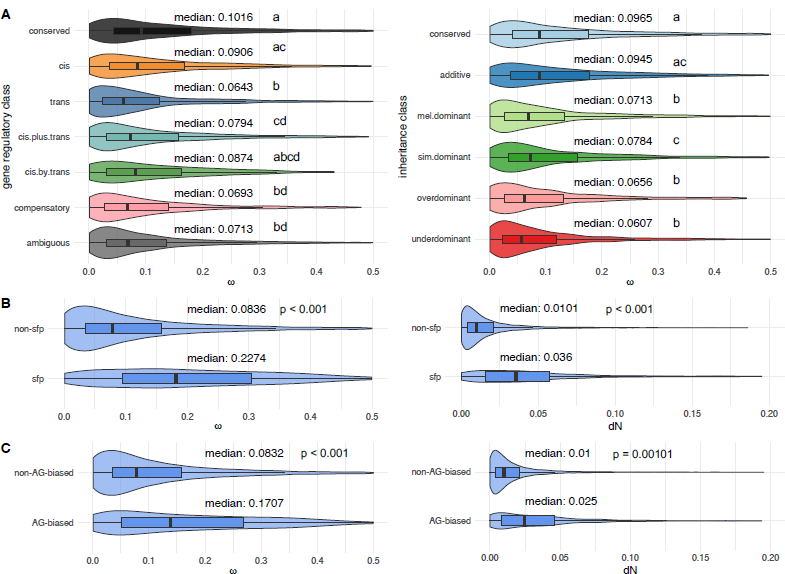
A) Distributions of ω among regulatory and inheritance classes. Alongside the median, significant differences by pairwise Wilcoxon rank sum tests (Holm-Bonferroni adjusted p < 0.05) are indicated by different numbers across gene sets. B) Distributions of dN and ω between Sfps and non-Sfps. C) Distributions of dN and ω between AG-biased and non-AG-biased genes. p values for B) and C) refer to Wilcoxon rank sum tests.

**Suplemental Figure 8.**
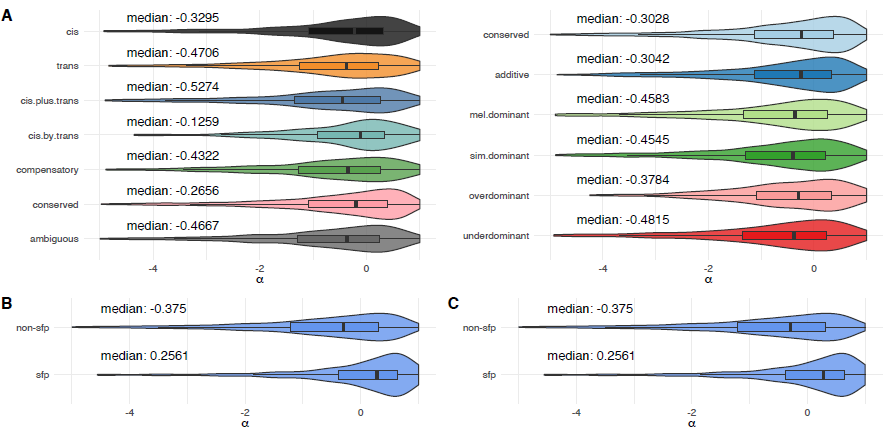
A) Distributions of ⍺ among regulatory and inheritance classes. Genes with conserved regulation have significantly greater median ⍺ than *trans* (Wilcoxon rank sum test, p = 03), *cis* + *trans* (p < 0.001), and ambiguous genes (p < 0.001); all other comparisons of inheritance s are not significantly different (p > 0.05). Genes with conserved inheritance have significantly greater median ⍺ than *mel* dominant (p = 0.001), *sim* dominant (p = 0.007), and underdominant genes (p = 0.021); all other comparisons of inheritance types are not significantly different (p > 0.05). B) Sfps e significantly greater median ⍺ than non-Sfps (p < 0.001). C) Distributions of ⍺ between AG-biased and non-AG-biased genes are not significantly different (p = 0.52).

**Supplemental Figure 9.**
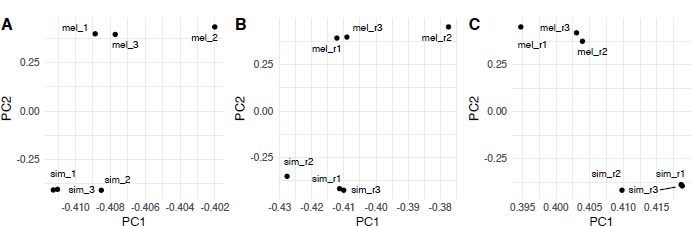
PCA of peak accessibility (log_2_ transformed counts) with the first two PCs shown: (A) conserved peaks, (B) *mel* orphan peaks, (C) *sim* orphan peaks.

**Supplemental Figure 10.**
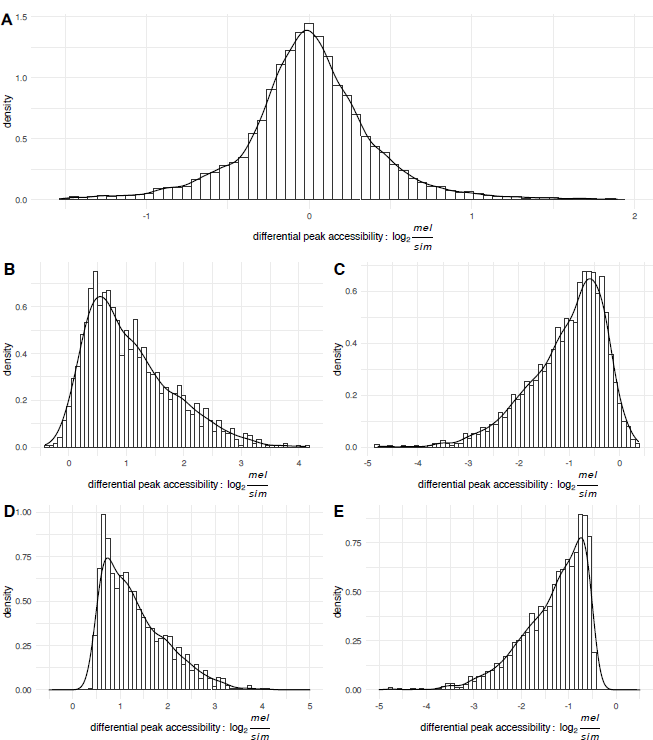
Differential accessibility (DA) of chromatin peaks called by ATAC-Seq, defined as the log_2_-transformed ratio of counts *D. melanogaster* counts to *D. simulans* counts. (A) Conserved peaks; (B) *mel* orphan peaks; (C) *sim* orphan peaks. We further filtered orphan peaks to just those that are DA and more highly expressed in the species with a peak called: (D) filtered *mel* orphan peaks; (E) filtered *sim* orphan peaks.

**Supplemental Figure 11.**
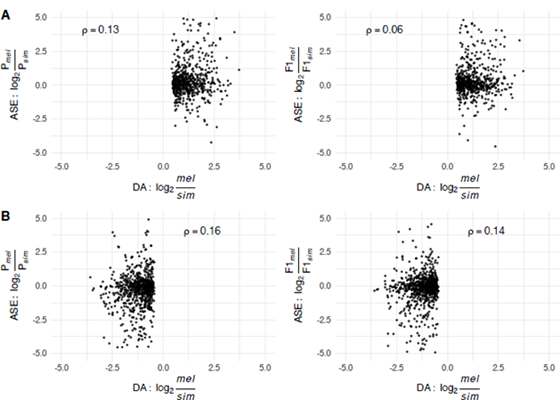
Parental expression divergence or ASE (y-axis) plotted against differential accessibility (DA) of chromatin peaks called by ATAC-Seq (x-axis). (A) *mel* orphan peaks; (B) *sim* orphan peaks. Spearman’s rank coefficient ⍴ is shown on each plot.

